# Bidirectional and context-dependent changes in theta and gamma oscillatory brain activity in noradrenergic cell-specific *Hypocretin/Orexin receptor 1*-KO mice

**DOI:** 10.1101/123075

**Authors:** Sha Li, Paul Franken, Anne Vassalli

## Abstract

Noradrenaline (NA) and hypocretins/orexins (HCRT), and their receptors, dynamically modulate the circuits that configure behavioral states, and their associated oscillatory activities. Salient stimuli activate spiking of locus coeruleus noradrenergic (NA^LC^) cells, inducing NA release and brain-wide noradrenergic signalling, thus resetting network activity, and mediating an orienting response. Hypothalamic HCRT neurons provide one of the densest input to NA^LC^ cells. To functionally address the HCRT-to-NA connection, we selectively disrupted the *Hcrtr1* gene in NA neurons, and analyzed resulting (*Hcrtr1^Dbh-CKO^*) mice’, and their control littermates’ electrocortical response in several contexts of enhanced arousal. Under enforced wakefulness (EW), or after cage change (CC), *Hcrtr1^Dbh-CKO^* mice exhibited a weakened ability to lower infra-θ frequencies (1-7 Hz), and mount a robust, narrow-bandwidth, high-frequency θ rhythm (~8.5 Hz). A fast-γ (55-80 Hz) response, whose dynamics closely parallelled θ, also diminished, while β/slow-γ activity (15-45 Hz) increased. Furthermore, EW-associated locomotion was lower. Surprisingly, nestbuilding-associated wakefulness, inversely, featured enhanced θ and fast-γ activities. Thus HCRT-to-NA signalling may fine-tune arousal, up in alarming conditions, and down during self-motivated, goal-driven behaviors. Lastly, slow-wave-sleep following EW and CC, but not nestbuilding, was severely deficient in slow-δ waves (0.75-2.25 Hz), suggesting that HCRT-to-NA signalling regulates the slow-δ rebound characterizing sleep after stress-associated arousal.

## Introduction

Electrical brain waves (or oscillations) can be used to define behavioral domains called vigilance (or behavioral) states. At any time, an animal appears to be in one of three ‘global brain states’, wakefulness, non-rapid-eye-movement (NREM) sleep (or slow-wave-sleep in rodents), and REM (or paradoxical) sleep, as assessed by the oscillatory profile measured in the electroencephalogram (EEG, or electrocorticogram (ECoG) in rodents). EEG signals provide information at a high-temporal resolution (10-100 ms) about an animal’s arousal state, behavior, and prior sleep/wake history. Behavioral states were also found to delineate distinct physiological and molecular brain milieus, e.g.^1,2^. Furthermore, increasing evidence suggests that oscillatory patterns are active functional determinants of brain circuits processing, and their behavioral outcomes, e.g.^3–6^.

State-specific oscillations emerge from dynamic neuronal activity in interconnected cortical and subcortical microcircuits responding to external stimuli and a multitude of internal indices of the animal’s circadian, homeostatic, and motivational state. Many of these signals are implemented by a set of key neuromodulators (acetylcholine, histamine, noradrenaline [NA], serotonin, dopamine, and hypocretin-1 and 2/orexin-A and B [HCRT]), produced in a set of subcortical arousal ‘hubs’, that are active when the animal is awake, but mostly silent in NREM sleep. Although all these neuromodulator-producing neurons regulate waking states and attention, they widely differ in their neuroanatomical and electrophysiological properties, the contexts that activate them, and the downstream effectors they act upon, and their behavioral translation^7,8^. The complex interplays between these neuromodulators are poorly understood.

HCRT and NA are produced by neurons residing in, respectively, the lateral hypothalamus and a set of pontomedullary NA nuclei^9^, of which the locus coeruleus (LC) is the best characterized. Each cell group receives multiple inputs and broadcasts to many nuclei, including to each other^10^. LC provides approximately half of all brain NA^11^, and all the neocortical and hippocampal NA. LC cells fire tonically at a frequency that is behavioral state-dependent, and in waking tends to increase with arousal level^12^. In response to salient stimuli, LC neurons shift to a 8-10 Hz phasic firing mode, which differentially regulates downstream circuit processing^13,14^. NA release in targets therefore depends on the firing pattern of NA^LC^ cells, which is modulated by many afferents. One of the densest innervation originates from HCRT neurons^9,15–17^. HCRT cell activity is enhanced by a wide range of behaviorally relevant stimuli, and was demonstrated to drive HCRT release in the LC, and to increase NA^LC^ neuron firing rate, both in slices^16,18,19^ and behaving animals^20^. HCRT in the LC, in turn was shown to evoke NA release in the dentate gyrus^21^, and the prefrontal cortex (PFC)^22,23^. How much of the LC response to salient cues reflects a primary response of HCRT neurons to these cues, remains unclear.

HCRT acts by signaling through two G-protein-coupled receptors, HCRTR1 and HCRTR2^24^, which are differentially expressed throughout the brain. *In situ* hybridization reveals that the LC, as well as the brainstem’s six other NA cell groups, express *Hcrtr1*, in relative absence of *Hcrtr2* expression^20,25,26^.

Optogenetic stimulation of both HCRT and NA^LC^ neurons increases the probability of sleep-to-wake transitions, but with much faster dynamics for NA^LC^, than HCRT neurons^27,28^. NA^LC^ optogenetic inhibition concerted with HCRT cell stimulation represses, while co-stimulation enhances, awakening^29^, establishing the HCRT-LC connection as a critical circuit to transition from sleep to wake. Loss of HCRT signaling by disruption of *Hcrt*, both *Hcrtr1* and *Hcrtr2* genes, or ablation of HCRT-producing cells, profoundly alters behavioral state regulation, and closely phenocopies narcolepsy symptomatology^30–32^. Loss of NA by disruption of the gene encoding the NA-synthesizing enzyme, *Dopamine-β-hydroxylase (Dbh*), produced contradicting reports, with indication of normal^33^, or reduced^34^ wakefulness in mice, and reduced wakefulness in zebrafish^35^.

Here we use genetic tools to inactivate the functional connectivity between HCRT neurons and downstream NA neurons. We generated a novel CRE recombinase-dependent *Hcrtr1* KO mouse strain, and we used a *Dbh-Cre* transgene^36^ to drive *Cre* expression in NA cells. In the resulting (*Hcrtr1^Dbh-CKO^*) mice, Dbh-expressing cells undergo Cre-mediated deletion of an essential *Hcrtr1* gene fragment, thus abolishing HCRTR1 signaling in all brain NA circuits. As HCRTR1 is thought to get transported to the NA cell membrane in the soma, the dendrites, and the axonal terminals, our mutant mice’ NA neurons lose the ability to respond both postsynaptically to HCRT (whithin NA cell groups), and presynaptically (within target tissues). Indeed NA release is finely regulated at the level of NA terminals, not only by autoinhibitory α2-adrenoceptors, but also by dopamine and other presynaptic receptors^37,38^. HCRTR1 was reported to act presynaptically at a variety of brain sites, regulating the release of glutamate, GABA or Ach^39–47^. Whether HCRTR1 acts presynaptically at NA terminals has, to our knowledge, not yet been investigated.

As HCRTR1 modulates behavioral arousal induced by both aversive and appetitive stimuli^8^, we assessed the global impact of the loss of HCRT-to-NA signaling by measuring brain oscillatory dynamics of *Hcrtr1^Dbh-CKO^* and control mice as they freely behaved in environments of both negative and positive emotional valence. Loss of HCRT-to-NA signaling was found to primarily affect the main ECoG correlates of arousal, i.e. θ and fast-γ frequencies. In stress-associated conditions, *Hcrtr1^Dbh-CKO^* mice’ waking exhibited increased δ and inter-δ/θ (4-7 Hz) activity, suggestive of reduced alertness, while normal induction of θ and fast-γ oscillations was blunted. Surprisingly, in spontaneous nocturnal nestbuilding-associated wake which precedes sleep, *Hcrtr1^Dbh-CKO^* mice displayed higher θ and fast-γ activity. Our study evidences the role of HCRT-to-NA signaling in building an appropriate θ/fast–γ response in stressing environments, and reveals circumstances in which HCRT-to-NA signaling may serve to curb hyperarousal.

## Results

### Generation of a Cre-dependent *Hcrtr1* KO mouse strain, and selective *Hcrtr1* gene inactivation in NA neurons

We modified *Hcrtr1* in embryonic stem cells so that the gene gets fully inactivated in presence of CRE, but remains functional in its absence (Fig. 1a). We inserted two loxP sites to flank an essential N-terminal-encoding region, and further modified the *Hcrtr1* locus so that CRE-mediated loxP site-specific recombination has three main effects: (1) it excises exons 3 and 4, which encode the signal and N-terminal polypeptide up to within transmembrane domain 3 (a total of 126 aa), (2) it replaces the *Hcrtr1* open reading frame with a (promoterless) *Gfp* open reading frame, allowing to use GFP expression to monitor cells which have undergone *Hcrtr1* gene disruption, but have an active *Hcrtr1* promoter, and thus would normally express *Hcrtr1*, and (3) it inserts a polyadenylation signal downstream of the GFP cassette, and thus terminates transcription and precludes expression of downstream exons. This conditional KO (CKO, or ‘floxed’) allele is referred to as *Hcrtr1^flox^*.

**Figure 1.**
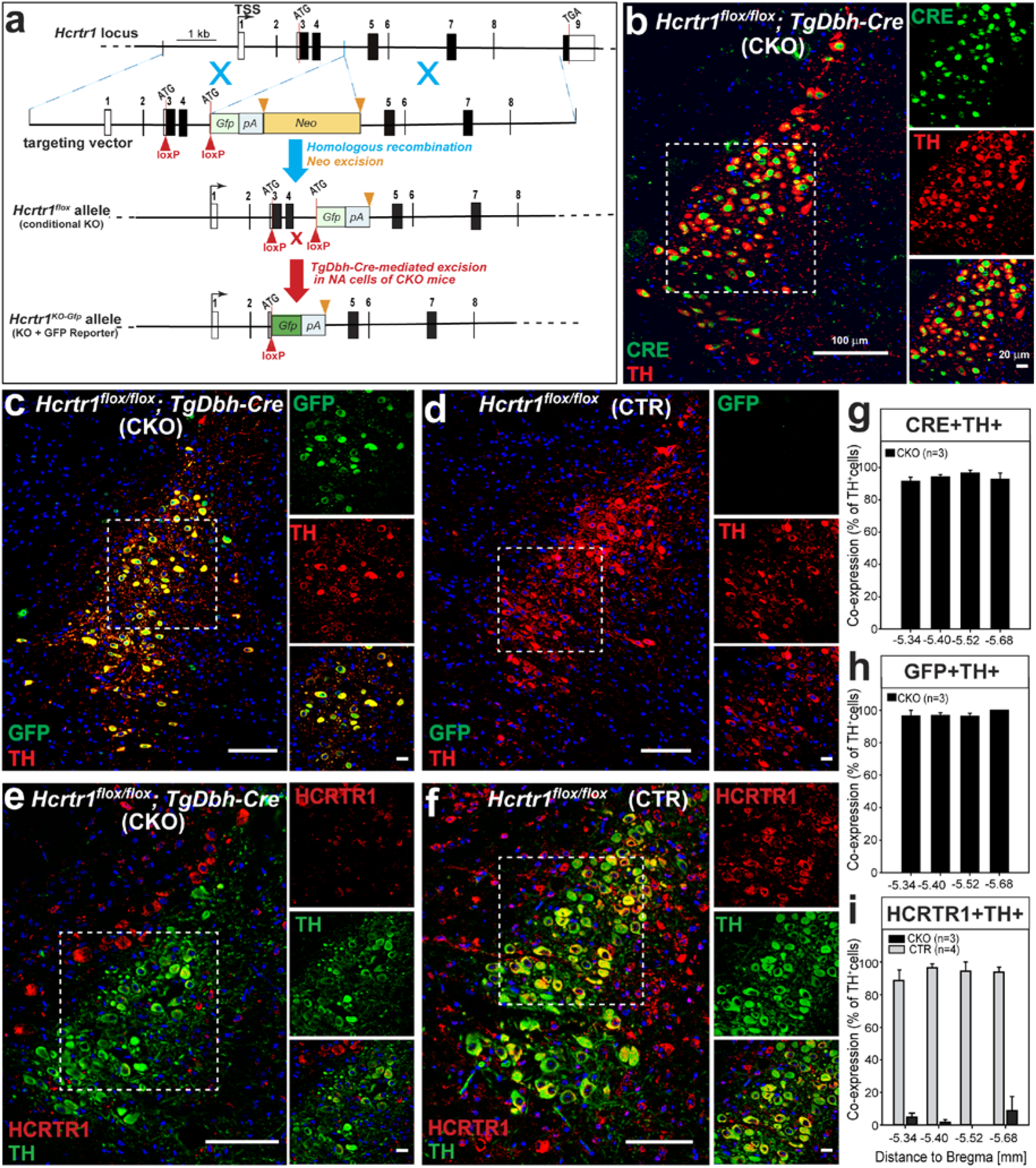
Conditional inactivation of *Hcrtr1* in noradrenaline (NA) neurons: TH^+^ cells in the locus coeruleus of *Hcrtr1^Dbh-CKO^* (CKO) mice express GFP in place of HCRTR1. (**a**) Schematic representation of homologous recombination between the *Hcrtr1* genomic locus and the targeting vector (TSS, transcription start site; filled rectangles depict protein-coding sequences, and unfilled rectangles represent untranslated regions). Two loxP sites (red triangles) were inserted to flank the first coding exon (exon 3) and exon 4, which together encode the N-terminal first 126 aa of HCRTR1. The neomycin resistance gene used for selection in embryonic stem cells (neo; flanked by two FRT sites shown as orange triangles) was deleted using the FLP recombinase, creating the *Hcrtr1^flox^* allele. In *TgDbh-Cre*-expressing cells, CRE excises the inter-loxP fragment (1.1 kb), creating the *Hcrtr_1_^+/KO-Gfp^* allele. In the latter, the translation start site (ATG) of *Hcrtr1* is replaced by the ATG of the *Gfp* cassette. Thus *Gfp* expression (green rectangle) becomes regulated by the *Hcrtr1* promoter. *Gfp* reading frame is followed by a polyadenylation site (pA) to terminate transcription and prevent downstream expression. (**b-f**) Representative confocal images of the locus coeruleus (LC) of mice of the indicated genotypes. Noradrenergic (NA) neurons are stained with a tyrosine hydroxylase antibody (TH^+^). Costaining with a CRE (**b**), GFP (**c-d**), or HCRTR1 (**e-f**) antibody is depicted. (**b**) shows that CRE is efficiently and specifically expressed in LC TH^+^ cells of a CKO mouse. (**c**) CKO mice express GFP in place of HCRTR1 in LC TH^+^ cells, while, in absence of CRE, (**d**) a CTR mouse lacks GFP immunoreactivity. (**f**) HCRTR1 immunoreactivity can be seen in LC TH^+^ cells of a CTR mouse, but not appreciably in LC TH^+^ cells of a CKO mouse (**e**). The CKO mouse exhibits however HCRTR1-immunoreactive, TH-negative cells outside the LC nucleus, consistently with NA cell-selective *Hcrtr1* gene disruption. The HCRTR1 antibody was raised against a peptide encoded by regions downstream of the CRE-mediated deletion. (**g-i**) Quantification of the percentage of TH^+^ neurons that are also immunoreactive for CRE (g), GFP (h), or HCRTR1 (i). Coronal sections collected at four levels throughout the LC were quantitatively assessed. Data are mean±SEM values for (g-h) CKO (n=3 for each), or (i) CKO (n=3) and CTR (n=4) mice. Boxed areas (and images to the right) represent fields delineated for cell counts (see Methods). The size indicated above scale bars in (b) applies also to (c-f). Mouse Genome Informatics: *Hcrtr1^flox^* is *Hcrtr1^tm1.1Ava^* (MGI:5637400), and *Hcrtr1^KO-Gfp^* is *Hcrtr1^tm1.2Ava^* (MGI: 5637401) (**www.informatics.jax.org/reference/j:226158**^50^).

To generate NA-specific *Hcrtr1* deficient mice, *Hcrtr1^flox/flox^* mice were crossed with mice harboring a *Dbh-Cre* BAC transgene^36^. We established breeding pairs that generate two offspring groups: *Hcrtr1^flox/flox^;TgDbh-Cre*-positive (*Hcrtr1^Dbh-CKO^*, or CKO) mice, and *Hcrtr1^flox/flox^;TgDbh-Cre*-negative (*Hcrtr1^Dbh-CTR^*, or CTR) mice. All analyses rely on pair-wise comparison between these two groups.

To demonstrate that accurate Cre-dependent DNA recombination occurs at the *Hcrtr1* locus, we prepared tissue punches from various tissues and brain areas from CKO and CTR mice, and the *Hcrtr1* gene was analyzed and sequenced. Only genomic DNA from the LC area of CKO mice amplified diagnostic recombined fragments of the *Hcrtr1* gene, whose sequencing confirmed accurate genomic structure and inter-loxP deletion breakpoints (see Methods).

### GFP labeling of NA^LC^ cells in *Hcrtr1^Dbh-CKO^*, but not *Hcrtr_1_^Dbh-CTR^* mice, evidences *Hcrtr1-> Gfp* open reading frame replacement

We first evaluated how efficient and specific the *Dbh-Cre* transgene used to target NA neurons was, by comparing tyrosine hydroxylase (TH) and CRE immunofluorescence in LC sections from CKO mice. Out of 732 TH-immunoreactive neurons, 689 expressed CRE (94.3±2.1%, mean±SEM for n=3 mice; Fig. 1b,g). Conversely, 96.8±1.4% of CRE-positive cells also expressed TH, confirming high penetrance and specificity of the transgene, respectively. Next, to determine efficiency and specificity of CRE-mediated recombination at the *Hcrtr1^flox^* locus, LC sections were assayed for TH and GFP co-expression. This revealed bright GFP immunoreactivity in NA^LC^ cells of CKO mice (Fig. 1c), and its absence in CTR mice (Fig. 1d). Out of 674 TH-positive cells in LC of CKO mice, 658 co-expressed GFP (97.6±1.3%, n=3; Fig. 1h), hence had undergone CRE/loxP recombination, while among 710 GFP-positive cells, 92.9±2.4% co-expressed TH, indicating, respectively, high penetrance and specificity of genomic deletion at the *Hcrtr1^flox^* locus.

### Loss of HCRTR1 immunoreactivity

To further demonstrate that genomic deletion precludes HCRTR1 protein expression, we assessed LC sections of CKO and CTR mice for TH and HCRTR1 immunofluorescence (Fig. 1e-f,i and see Methods). Among 523 TH-positive cells in the LC of CTR mice, 498 were positive for HCRTR1 (93.3±3.7%, mean±SEM for n=4 mice; Fig. 1f,i), while among 431 TH-positive cells in LC of CKO mice, only 11 were HCRTR1-positive (2.8±2.0%, n=3; Fig. 1e,i). Consistently with the NA-cell specificity of *Hcrtr1* disruption, HCRTR1-immunoreactive, TH-negative cells were present in brain areas outside the LC (Fig. 1e). These data demonstrate highly penetrant, Cre-dependent *Hcrtr1* KO in the LC TH+ neurons of our *CKO* mouse model.

To further validate our *Hcrtr1* mutagenesis strategy, we generated mice carrying the recombined (*KO-Gfp) Hcrtr1* allele (Fig. 1a) in all cells. To this end we crossed *Hcrtr1^flox/flox^* animals to EIIa-Cre transgenic mice^49^, a line expressing CRE in the pre-implantation embryo (see Methods). Early embryonic CRE/loxP recombination allowed transmission of the recombined *Hcrtr1* gene into the germ-line, creating a *Hcrtr1^KO-Gfp^* mouse strain. Homozygosity of the mutation is expected to provide a *Hcrtr1* null. Furthermore, since *Gfp* is under control of the endogenous *Hcrtr1* promoter, heterozygous *Hcrtr1^+/KO-Gfp^* animals permit mapping of *Hcrtr1* gene activity across the organism^50^. As expected, nearly all TH+ cells in the LC of *Hcrtr1^+/KO-Gfp^* mice express GFP (see Supplementary Fig. S1a online). Out of 709 TH-expressing cells (n=3 mice), 99.6±0.3% also expressed GFP, and 91.2±4.6% of GFP-positive cells in the LC also expressed TH, confirming the robust activity of the *Hcrtr1* promoter in NA^LC^ cells.

We next analyzed HCRTR1 immunofluorescence in mice heterozygous (n=3), or homozygous (n=3) for the *Hcrtr1^KO-Gfp^* mutation. LC imaging confirmed loss of HCRTR1 immunofluorescence in homozygous *Hcrtr1^KO-Gfp/KO-Gfp^* mice (Fig. S1b, c). In contrast to CKO mice, the latter mice have lost HCRTR1-immunoreactivity not only in LC TH+ cells, but in areas surrounding the LC as well (Fig. S1c). These data support the specificity of the antibody used in Fig. 1e-f to assess CKO and CTR mice, and creation of a constitutive *Hcrtr1* KO (null) mouse model.

### Increased δ oscillatory activity in wakefulness of *Hcrtr1^Dbh-CKO^* mice

A cohort of CKO (n=9) and CTR (n=7) animals were submitted to several behavioral paradigms across a 9-day period (Fig. 2) as their ECoG/EMG signals were recorded. In undisturbed conditions (baseline), total time spent in wakefulness (W), slow-wave-sleep (SWS), or paradoxical sleep (PS), and their temporal dynamics over the 24-h day did not differ between the two genotypes (Figs 3a, S2, S6, and Supplementary Table S1). To assess the finer architecture of sleep-wake states, we quantified the distribution of episode duration, both with respect to episode number as well as the percentage of total time in each state spent in episodes of various durations (Figs S2-S3). Sleep-wake architecture was very similar for the two genotypes with the exception of PS. For this state we observed a right shift of the distribution of percent time spent in PS across episode duration categories (Fig. S2b *Right;* interaction for factors ‘episode duration’ and ‘genotype’ two-way ANOVA, *P*<0.002), with CKO mice spending most of PS in ~2-min bouts (64-128 s), while CTR mice spent most of PS in ~1-min bouts (32-64 s).

**Figure 2.**
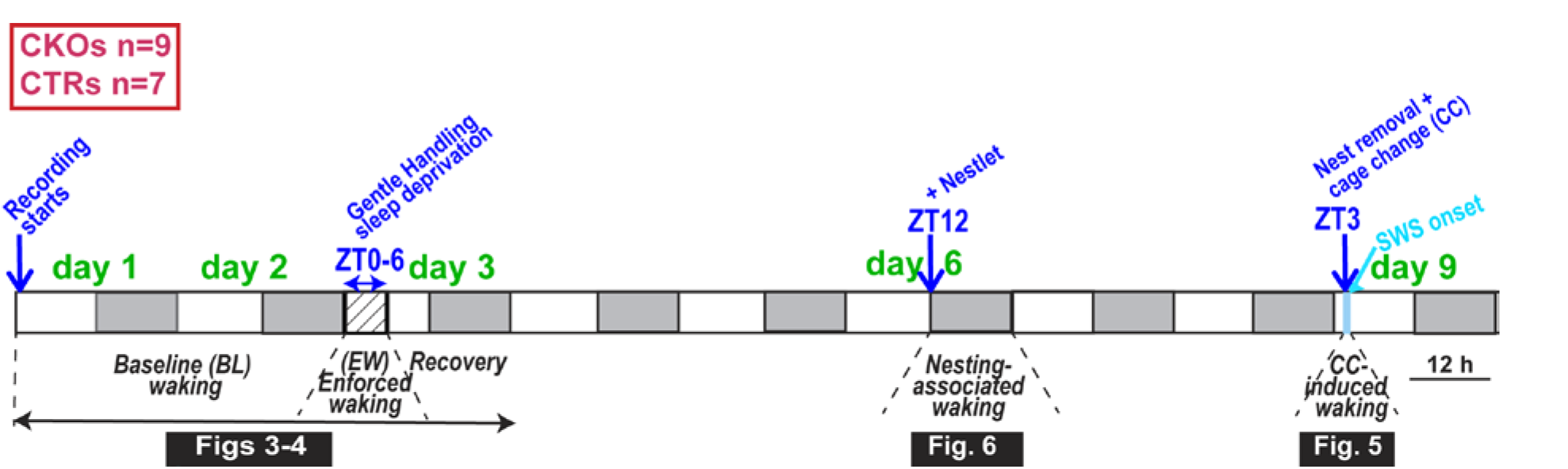
Experimental design. A cohort of *Hcrtr1^Dbh-CKO^* (n=9) and *Hcrtr1^Dbh-CTR^* (n=7) littermate mice were sequentially exposed to distinct behavioral paradigms, as their ECoG was recorded. Spontaneous waking ECoG activity was analyzed in the first two baseline (BL) days. Starting at light onset (ZT0) of day 3, mice were submitted to total sleep deprivation for 6 hours (6h) in their home cage (EW or enforced waking). Mice were then left undisturbed for two days. At dark onset (ZT12) of day6, nest material was introduced (+Nestlet), and the waking ECoG analyzed in the following 24h. After an additional two intervening days, 3h after light onset (ZT3) of day 9, each mouse was removed from its nesting cage, and transferred to a fresh cage (cage change, CC). CC-induced waking was analyzed until the next SWS-onset. Shaded areas represent the 12h dark phases.

**Figure 3.**
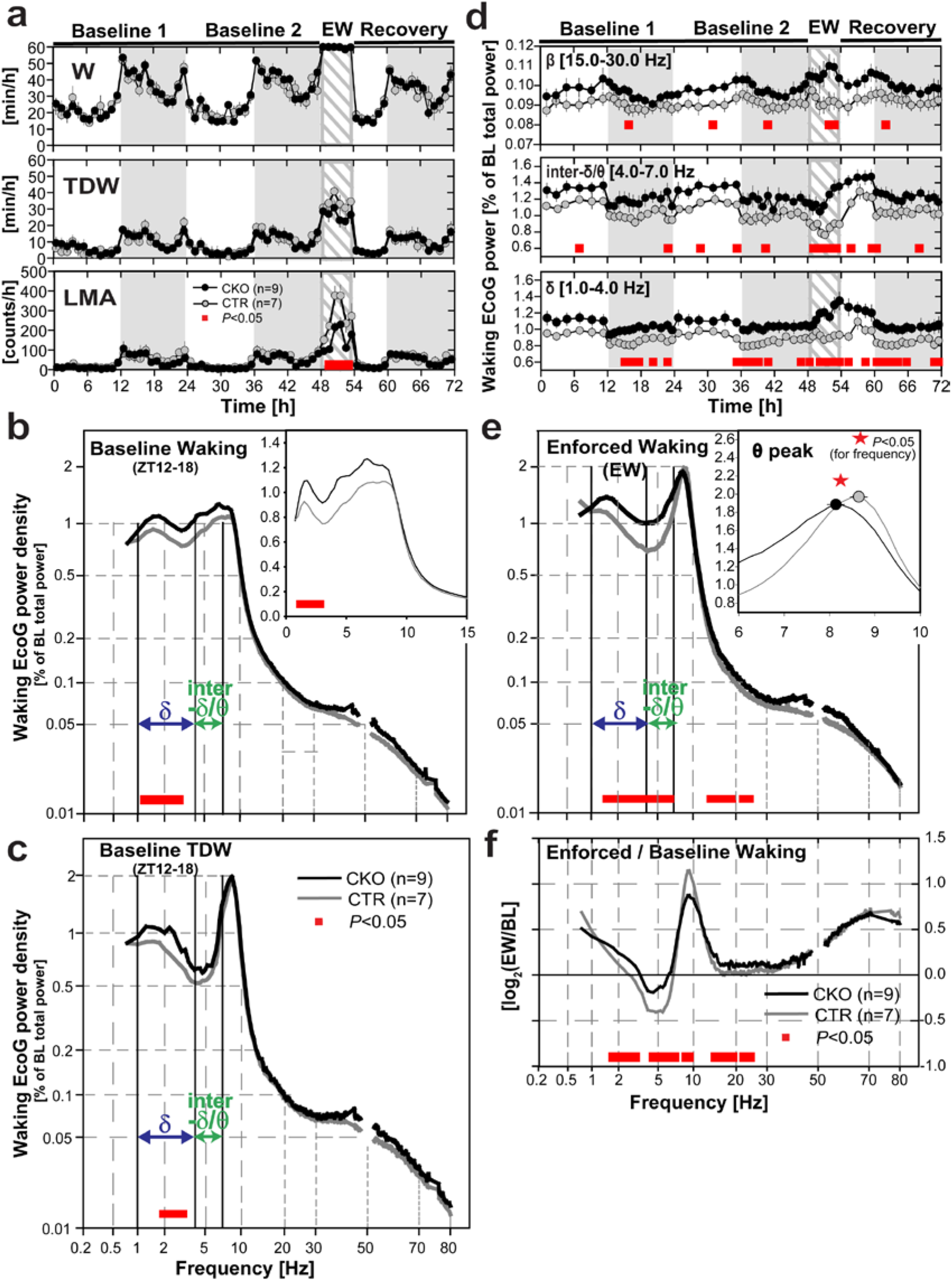
Spontaneous vs. enforced wakefulness. (**a**) Mean (±SEM) hourly values of (*Top*) total waking time [min/h], (*Middle*) theta-dominated-waking (TDW) [min/h], and (*Bottom*) locomotor activity (LMA) [counts/h] across experimental days 1-3 in *Hcrtr1^Dbh-CKO^* (CKO, ●), and *Hcrtr1^Dbh-CTR^* (CTR, ○) mice. ECoG spectra are shown for: (**b**) wakefulness in baseline dark phase ZT12-18, (**c**) TDW in baseline dark phase ZT12-18, and (**e**) enforced waking ZT0-6. ECoG power density is expressed as % of a baseline power reference value, calculated for each mouse across all states and all frequencies over 48h^134^(see Methods) [% of ‘BL total power’]. Note non-linear axes. Insets in (b) and (e) show ECoG spectra magnified across frequency ranges of interest, depicted with linear axes. Waking spectra of CKO mice differed significantly from those of CTR mice in all 3 cases (b-c, and e). Spectra depicted in b and e demonstrate significant genotype, and genotype X frequency effects across 0.75-80 Hz (two-way ANOVA, *P*<0.001 for all). Baseline TDW (c) demonstrates a significant genotype effect (two-way ANOVA, *P*<0.001), with no genotype X frequency interaction (*P*=0.249). CKO mice’ baseline waking showed enrichment in frequencies in the δ range (1.25 to 2.75 Hz) relative to CTR littermates (b). During EW (e), CKO’ ECoG power increase extended to encompass the 1.5-6.75 Hz (δ + inter-δ/θ), and 13.5-24 Hz (β) frequency ranges (post-hoc independent two-tailed student *t*-tests, *P*<0.05). Inset in e shows the theta rhythm’s peak frequency (TPF) during EW in CKO (●), and CTR (○) mice (mean±SEM). TPF is slower in CKO mice (^✱^, two-tailed t-test, *P*<0.05). (**f**) To extract the spectral traits specific to the EW response within each genotype, EW power spectra are expressed relative to baseline waking power spectra (as averaged across 48h in recording days 1-2). EW/baseline power density ratio is expressed as log_2_ with 0 indicating no change and ±1.0 indicating a 2-fold change. CKO mice’ EW-response is marked by a lesser increase in the θ range (bins concerned: 8.75-9.5 Hz) and a lesser decrease in the δ and inter-δ/θ ranges (bins concerned 1.75-3 Hz and 4.5-7 Hz). (**d**) Timecourses of waking δ (1-4 Hz), inter-δ/θ (4-7 Hz), and β (15-30 Hz) activity during days 1-3, expressed as % of the same baseline total ECoG power reference as used for the waking spectra in b-c and e. Red bars indicate significant genotype differences (post-hoc independent two-tailed student *t*-tests that followed two-way ANOVAs, *P*<0.05).

Furthermore, during the 18h for which recovery sleep was monitored after the 6-h sleep deprivation, hourly sleep-wake state amounts did not differ between the two genotypes (Figs 3a and S6), again with the exception of PS (Fig. S2a). The timecourse of hourly values of PS time was significantly altered by genotype (interaction factors ‘time’ and ‘genotype’ two-way ANOVA, P=0.01). CKO mice spent significantly more time in PS than did CTR mice in the middle of the recovery light period [ZT8-10], which is the period of maximal PS rebound for both genotypes. In that interval, CKO expressed a 26% increase in PS time over CTR mice (CKO: 14.4±0.5 min vs CTR: 11.4±0.4 min; t-test p=0.0005), which was due to a 20-s lengthening of mean PS episode duration (+31%) (CKO: 84.7±5.6 s vs CTR: 64.9±4.1 s; t-test, P=0.018), leaving episode number unaffected (−5%; t-test, p=0.691). Altogether these data suggest that PS maintenance, but not initiation, is favoured in CKO mice. An increase in PS state stability in CKO mice may be related to the known PS-suppressing activity of the HCRT-1 peptide, as demonstrated when locally applied in LC of rats^20^.

While the duration of CKO mice’ baseline waking was unchanged, its spectral quality was. Spontaneous wakefulness of CKO mice showed increased ECoG activity within the δ frequency-range (1.25 to 2.75 Hz) relative to CTR littermates in the dark phase first 6h (ZT12-18) (Fig. 3b). To delineate the times of day in which CKO mice’ wake differs in spectral activity, the power dynamics of the waking ECoG in specific frequency bands were plotted across days 1-3 using the same power reference as used in the waking spectra of Fig. 3b (see Methods). This revealed that CKO mice’ waking state is enriched in δ frequencies throughout recording days 1-3, but particularly at times of increased behavioral activity (i.e., in dark phase and during EW; Fig. 3d Bottom). It thus appears that at critical times of enhanced arousal, CKO mice are impaired in reducing δ activity as normal mice do. Heightened δ activity while awake is associated with drowsiness and increase sleep pressure in rodents and humana^51,52^.

We then asked whether the enhanced waking δ activity of CKO mice could also be observed during active waking, or more specifically in the ‘theta-dominated waking substate’ (TDW), we previously formulated an ECoG-based algorithm to identify^53^. We found that CKO mice’ baseline TDW state features increased power in the δ range (2.0-3.0 Hz) relative to CTR mice as well (Fig. 3c). Further analysis revealed that increased δ power also affects CKO mice’ non-TDW waking (akin to ‘quiet waking’, not shown). Therefore the weakened ability of CKO mice to downregulate δ oscillations across waking substates suggests they suffer a globally reduced level of alertness.

### Enhanced infra-θ oscillatory activity, and impaired θ and fast-γ power induction in enforced wakefulness of *Hcrtr1^Dbh-CKO^* mice

At light onset (ZT0) of day 3, when sleep propensity is maximal, an experimenter entered the room, and the gentle handling sleep deprivation procedure was initiated and maintained for 6h. CKO mice’ ECoG spectra differed more extensively from CTR mice spectra in EW than they did in baseline waking (Fig. 3e). ECoG power across 1.5 to 6.75 Hz, encompassing δ and some of the traditionally defined θ range, and across 13.5 to 24 Hz (β) frequencies were enhanced in CKO mice. A prominent component of this activity increase occurs in the 4.0-7.0 Hz window, i.e., above the δ range but below the θ rhythm characteristic of explorative locomotion and voluntary behaviors in rodents^54^. We refer to the 4.0-7.0 Hz range in mice as the ‘inter-δ/θ’ band (although it is sometimes included within the ‘θ’ window by others^55^. We reserve ‘θ ‘ for the higher-frequency (7.5-11.5 Hz), narrow-bandwidth, hippocampal oscillations that are specifically induced upon active waking, and define the TDW state^53^. In contrast, the 4.0-7.0 Hz activity (see peak in baseline waking ECoG spectra, Fig. 3b) dominates during quiet waking and automatic behaviors, and it increases as sleep homeostatic pressure rises^51,55^.

Next, we examined the dynamics of the two other oscillatory ranges associated with reduced arousal (inter-δ/θ, and β) (Fig. 3d). Throughout the three days, CKO mice exhibited higher δ, but also inter-δ/θ, and, to a lesser degree, β (15-30 Hz) activity, compared to CTRs. Interestingly, CKO mice exhibited a steep power increase in all three bands in the course of EW, while this was not the case in control mice. This again indicates that CKO mice have an impaired ability to repress infra-θ oscillations at times of enhanced arousal, and furthermore suggests a faster build up rate of electrocortical signs of sleep pressure.

In both CKO and CTR mice, EW is singularized from average baseline waking by emergence of a sharp θ rhythm (Figs 3e and 4a). EW in CKO mice however differed from CTR mice’ EW in θ frequency and power. In CKO mice, the θ rhythm was significantly slower (Fig. 3e inset; CKO vs. CTR, 8.1±0.1 Hz vs. 8.6±0.1 Hz; *t*-test; t_14_ =-3.749; *P*=0.0022; and see Supplementary Table S2). To extract the spectral features of each genotype’s EW-response, EW spectra were contrasted to baseline waking spectra (Fig. 3f). This revealed that the increased θ, and decreased inter-δ/θ activity, that characterize the CTR mice’ response during the gentle-handling procedure, were significantly diminished in CKO mice. Lower activity in the θ range, but higher infra-θ (δ and inter-δ/θ) activity mark the blunted EW θ response of CKO mice, and both are suggestive of a weaker medial septal/hippocampal activation^56^.

**Figure 4.**
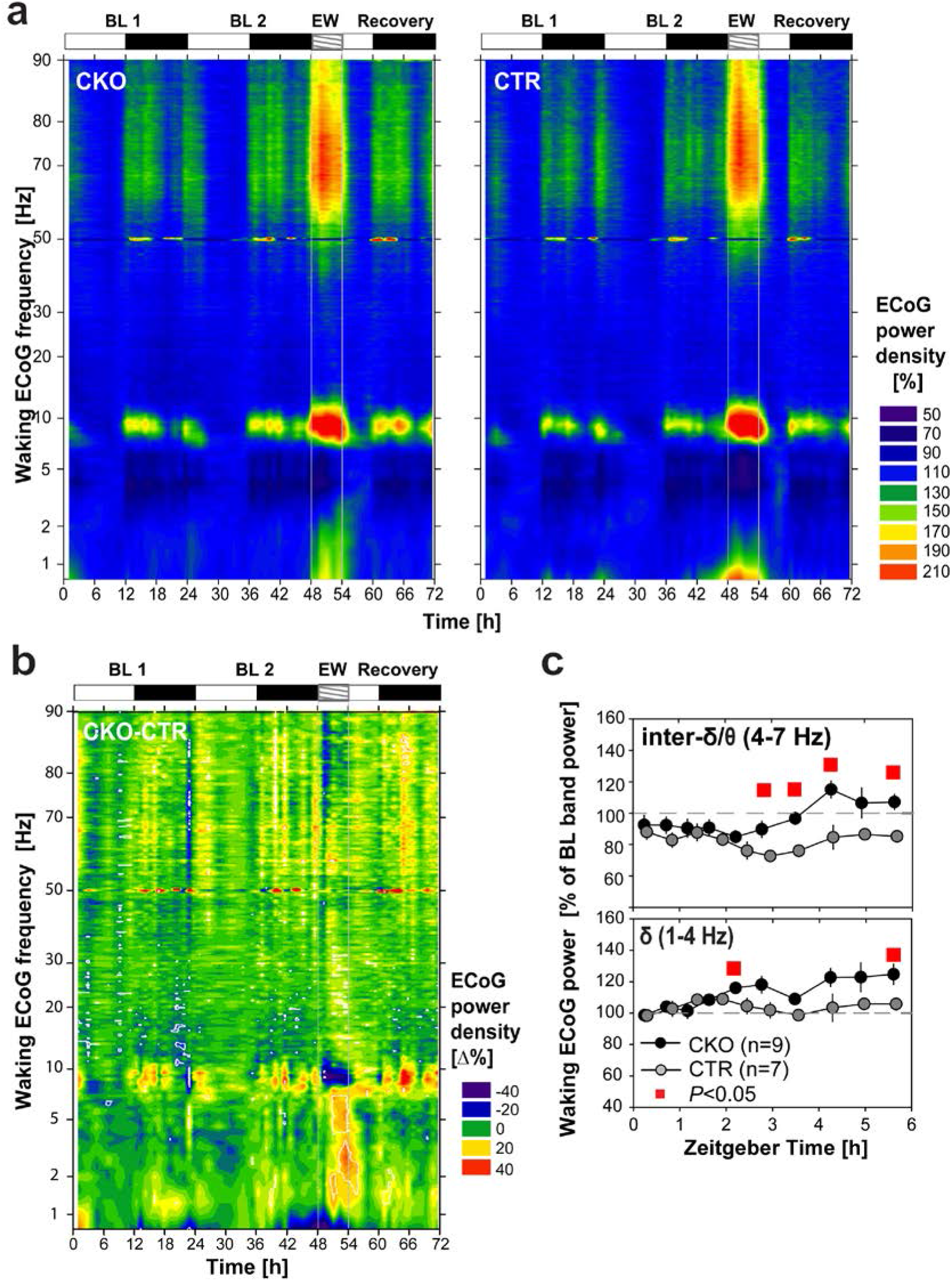
*Hcrtr1^Dbh-CKO^* mice exhibit weakened θ and fast-γ activity responses during gentle handling-enforced wakefulness (EW). (**a**) Heatmap depicting the ECoG spectral dynamics of the waking state across days 1-3. ECoG power in each 0.25 Hz frequency bin is expressed as % of its average value in the last 4h of the light phase (ZT8-12) in baseline days 1-2. Values are color-coded according to the scale shown on the right. (**b**) Differential dynamics of the waking ECoG of CKO and CTR mice (CKO-CTR). Spectral dynamics shown in (a) were substracted from each other [Δ%]. While both CKO and CTR mice’ waking state exhibit sharp increases in θ and fast-γ activity during EW (a), CKO mice’ waking exhibits a markedly lower θ, as well as fast-γ power, relative to CTR mice during EW, in particular in the first hour (dark blue), and a concomitant increase in infra-θ power. Episodes in which CKO-CTR differential activity shows, inversely, higher θ and fast-γ, power in CKO, occur during spontaneous waking, in particular in the dark phase, at times of high wakefulness (see Fig. 3a), during both baseline and recovery night. Such particularly pronounced episodes appear to occur during the recovery night after EW. White lines delimit areas in which power density significantly differs between genotypes (two-tailed t-tests, *P*=0.05). Note that color scales in (a) and (b) differ. (**c**) Timecourses of the waking ECoG activity in the δ (1-4 Hz) and inter-δ/θ (4-7 Hz) frequency bands across 6-h of EW, are depicted, with power expressed, as in (a), as % of its mean value in baseline ZT8-12 wakefulness.

Robust θ oscillations and TDW expression characterize vigorous explorative locomotor activity (LMA), but can also uncouple from it^53^, and be observed in immobile rats following a running bout^57^, or in association with motivational activation^58^. To investigate this relation in our mutant, LMA was monitored. While LMA was normal throughout the two baseline days, its increase during the 6-h EW period was markedly reduced in CKO mice (Fig. 3a Bottom; CKO vs. CTR, 929±120 vs. 1631±190 counts; Mann-Whitney U test, *P*=0.01). We previously found that mice lacking all HCRT signalling (*Hcrt^ko/ko^*) exhibit reduced locomotion relative to WT littermates throughout baseline and EW^53^, with a reduction during EW that was very similar in extent to the one we observe here in CKO mice relative to their CTR littermates. This suggests that the role of HCRT in locomotion relies primarily on the HCRT-to-NA pathway in EW, but on other HCRT targets in spontaneous waking.

To investigate more globally the spectral dynamics of wakefulness, an ECoG power heatmap was generated as a function of frequency and time, across the entire 0.75-90 Hz spectral range, and days 1-3 (Fig. 4a). As waking spectra show a strong dependence on prior sleep/wake history and homeostatic sleep pressure, the waking ECoG was referenced to baseline light phase last 4 h, i.e., after the major sleep period, when sleep pressure has returned to trough values (see Methods, Fig. 4 legend, and Fig. S4). This analysis revealed in both CKO and CTR mice powerful θ and fast-γ activity surges occurring in concert during EW, as well as, but to a lesser degree, in baseline waking of the first half of the dark period (Fig. 4a).

To next specifically delineate the genotype-based differences in this response, the two spectral dynamics obtained in Fig. 4a were subtracted from each other to produce the CKO-CTR differential profile in waking ECoG dynamics shown in Fig. 4b. This reveals a number of genotype differences. *CKO* mice feature a prominent deficit in enhancing θ oscillations during EW (blue color-code in b). θ oscillatory power induction, bandwidth narrowing, and frequency increase that characterized CTR EW are all weakened in CKO mice. Furthermore, a 70-90 Hz activity, closely paralleling θ power dynamics, was found to surge during EW, and to be similarly weakened in CKO mice. This is in line with the reduced θ power response of CKO mice in EW, as observed by spectral analysis in Fig. 3f. Surprisingly, some time intervals in the active (dark) phase, conversely, appeared to be associated with enhanced θ activity in CKO relative to CTR mice (Fig. 4b), a finding further replicated, and extensively discussed, below. Analysis of the dynamics of δ (1-4 Hz) and inter-δ/θ band (4-7 Hz) oscillatory power throughout EW 6-h revealed enhanced δ, and inter-δ/θ activity in CKO mice, in particular in EW’ 2nd half (Fig. 4c), a finding consistent with EW spectral profiles (Fig. 3e-f). Altogether our data suggest that the hippocampal θ and fast-γ oscillatory responses to EW are blunted in CKO mice. Hence we propose that HCRTR1-dependent neuromodulation of NA microcircuits contribute significantly to these responses.

### *Hcrtr1^Dbh-CKO^* mice exhibit weakened θ and fast-γ oscillatory responses to cage change

Following EW and recovery period, mice were left undisturbed for 2 days. At dark onset of day6, each mouse was provided nesting material, and left undisturbed for another 2 days (see next section). To assay capacity to adapt to a new environment, mice were transferred from their nesting home cage to a clean cage at ZT3 (see Fig. 2), a time ordinarily spent predominantly asleep (Fig. S6). Cage change (CC) had a powerful arousing effect lasting between 1.5-2h in both genotypes, (Fig. 5a*Top*). The ECoG response to CC however differed between the two groups (Fig. 5c). An active waking-like θ activity peak emerges in both mouse groups, but it appears blunted in CKO mice.

**Figure 5.**
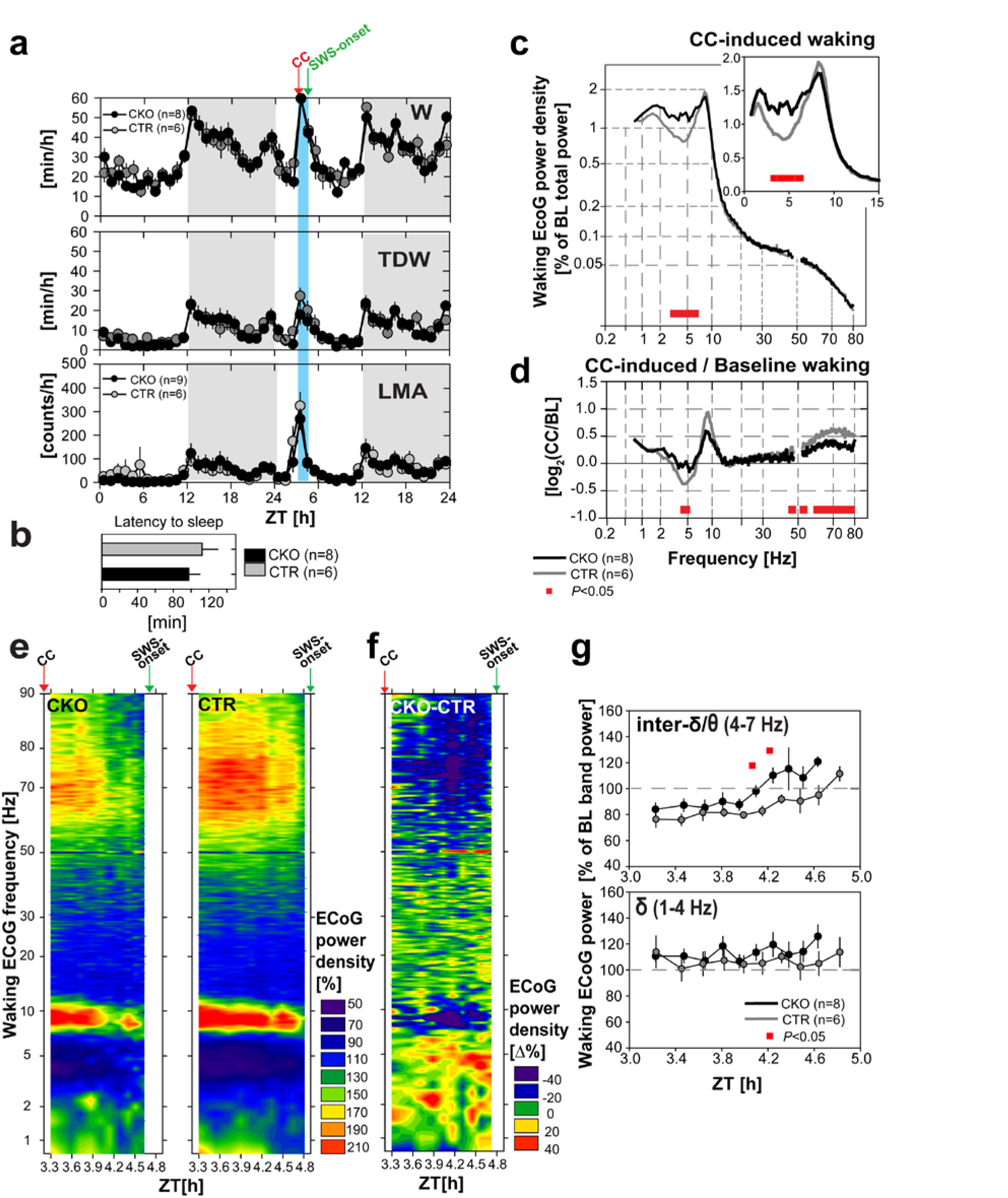
*Hcrtr1^Dbh-CKO^* (CKO) mice display weakened θ and fast-γ oscillatory responses upon transfer from their nesting cage to a clean cage. Mice were transferred to a fresh cage at ZT3 of the 9^th^ recording day. (**a**) Mean (±SEM) hourly values of (*Top*) total waking time [min/h], (*Middle*) theta-dominated-waking (TDW) [min/h], and (*Bottom*) locomotor activity (LMA) [counts/h] in the day preceding, and the day in which cage change (CC) occurred in CKO (●) and CTR (○) mice. (**b**) Latencies to SWS-onset are calculated as the time between CC and the first SWS episode [minute]. (**c**) Waking ECoG spectra from CC to SWS-onset in *Hcrtr1^Dbh-CKO^* (CKO, n=8), and *Hcrtr1^Dbh-CTR^* (CTR, n=6) mice, with power density values, expressed as % of each animal’s baseline ECoG ‘total power’ reference, as for spectra shown in Fig. 3b, c and e. Note the non-linear axes. CC-induced waking spectra of CKO and CTR mice differed, demonstrating significant genotype, and genotype X frequency effects across 0.75-80 Hz (two-way ANOVA, *P*<0.001 for all). CC induces a θ oscillatory peak in both genotypes, but its shape appears blunted in CKO mice. (**d**) To next extract CKO and CTR mice’ CC-induced changes in spectral oscillatory activity, CC-induced power spectra (as shown in c) were expressed relative to the mice’ baseline waking spectra (as averaged across the preceding 24h period) (CC-induced/Baseline waking). Power density ratio is expressed as log_2_. CKO mice exhibit a lesser decrease in inter-δ/θ band activity, and a profoundly diminished 53-80 Hz (fast-γ) response compared to controls (two way ANOVA indicated significant genotype and genotype X frequency effects, *P* <0.001 for both). Red bars indicate significant genotype differences (post-hoc independent two-tailed *t*-test). (**e**). Spectral dynamics of the waking ECoG after cage change, across time (from ZT3 to SWS-onset), frequencies (0.75 -90.0 Hz), and power (expressed as % of each mouse’ mean waking ECoG power in that frequency bin in the preceding day’s ZT8-12 time interval; color-coded as shown on the right. x-axis shows ZT time, adjusted to individual waking latencies to SWS-onset. (**f**) The differential CKO-CTR waking power dynamics following CC are depicted as they were for EW (Fig. 4b). Note that color codes for (e) and (f) differ. (**g**) Timecourses of ECoG power in two infra-θ frequency bands (1-4 Hz; 4-7 Hz) that were seen to respond in CC-induced waking in (e-f). ECoG power is expressed relative to the power in that band in average preceding day wakefulness. □, time intervals in which ECoG power significantly differed between CKO and CTR mice (independent two-tailed student *t*-test, *P*<0.05).

When waking spectra were analyzed specifically for their CC-induced features, (by plotting power spectral density values following CC relative to each group’s own average baseline values (Fig. 5d), and CC responses were contrasted between the two genotypes, CKO mice were seen to exhibit (i) a lesser CC-induced reduction in inter-δ/θ activity, similar to what was observed in EW (Fig. 3f), and (ii) a profoundly diminished fast-γ response across 55 to 80 Hz.

Heatmap representations of the spectral dynamics of the waking state expressed from CC to the subsequent SWS-onset (Fig. 5e), and the differential CKO-CTR dynamics thereof (Fig. 5f), confirm the weakened ECoG response of CKO mice. They powerfully illustrate CKO mice’ blunted θ rhythm response, compared to CTR mice’ response, as well as CKO mice’ impaired inter-δ/θ band activity dampening relative to its value in preceding light phase last 4h (Fig. 5e left, and 5f), while this CC-induced dampening is evident in controls (Fig. 5e right). CKO mice’ weakened fast-γ response is also clearly apparent (Fig. 5e-f).

Analysis of the dynamics of δ (1-4 Hz) and inter-δ/θ band (4-7 Hz) oscillatory power in CC-induced waking (Fig. 5g) revealed a rise in inter-δ/θ activity in both genotypes as time progresses. This increase was steeper however in CKOs, in particular in the 2^nd^ hour (independent two-tailed student’s *t*-test, *P* <0.05). In sum, although CC enhances θ synchrony and fast-γ oscillatory power in both CKO and CTR mice, weakening of both processes in CKO mice suggests HCRT-dependent noradrenergic activation participates in enhancing θ/fast-γ network activity in response to adverse environmental changes.

### Inverse θ and fast-γ dynamics of *Hcrtr1^Dbh-CKO^* mice in nesting-associated waking compared to enforced, or CC-induced, waking

To next explore our mice’ response to a novel, but unthreatening behavioral context, animals were recorded in a nestbuilding paradigm. Nesting material is rewarding in laboratory mice, and induces nestbuilding, a remarkably robust behavior^59,60^. A square of shreddable compressed cotton adapted for mouse nestbuilding (Nestlet^™^), which the mice had never experienced before, was introduced in the homecage at dark onset (ZT12) of day 6 (see Fig. 2). Nestlet-to-nest conversion was confirmed at next light onset in all animals (CKO n=5; CTR n=5) (see Methods).

Video-based behavioral analysis of a subset of mice (CKO n=3; CTR n=1) evidenced two major nest material interactive periods during the night: an early-night ‘discovery phase’, with numerous, brief episodes in which mice explored/shortly manipulated the Nestlet, and a late-night phase with longer manipulating bouts, and sustained nestbuilding (Fig. S5). Late-night bouts typically preceded SWS episodes. CKO mice spent more time awake than CTR mice in the time interval immediately preceding light onset (Fig. 6a).

**Figure 6.**
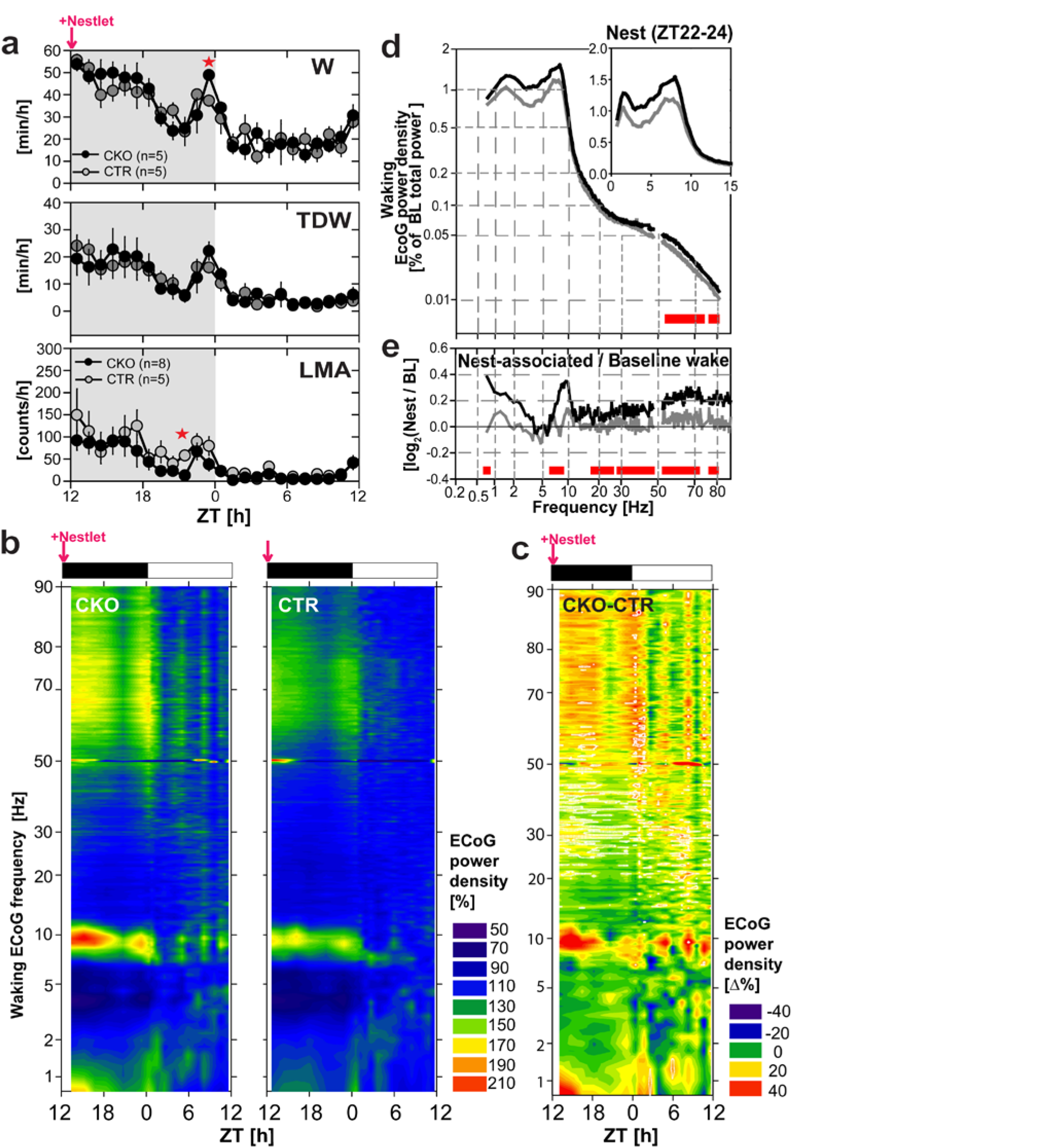
Availability of nesting material induces a stronger θ and fast-γ oscillatory response in *Hcrtr1^Dbh-CKO^* (CKO) mice than it does in control littermates in late night wakefulness that precedes sleep. Nest material (a Nestlet^TM^) was provided at dark onset (ZT12), and the ECoG was recorded in the following 24h. (**a**) Mean (±SEM) hourly values of (*Top*) total waking time [min/h], (*Middle*) theta-dominated-waking (TDW) [min/h], and (*Bottom*) locomotor activity (LMA) [counts/h] across the night that followed Nestlet addition, and the following light phase. ^✱^ denote time intervals with significant genotype differences (*t*-test, *P*<0.05) (b) ECoG power heatmaps depict the spectral dynamics of the waking state across the nesting night and the following light phase as shown in Fig. 4a. Plotted are ECoG power densities for each 0.25 Hz frequency bin relative to their values in baseline wakefulness across the ZT8-12 period [%]. (**c**) Differential CKO-CTR heatmap of the nesting 24h period was constructed similarly to Fig. 4b. Note that the color code for (e) and (g) differ. (**d**) Waking ECoG spectra in the last 2h (ZT22-24) of the nesting night are plotted as % of the baseline total power reference [%], as in Fig. 3b-c, e. CKO and CTR mice’ spectra demonstrate significant genotype, and genotype X frequency effects across 0.75-80 Hz (two-way ANOVA, *P*<0.001 for all). (**e**) ZT22-24 waking spectra shown in (d) are contrasted to waking spectra at the same time of day in baseline. The resulting Nesting/Baseline waking ECoG power density ratio demonstrates a significant genotype effect (two-way ANOVA, *P*<0.001), but no genotype X frequency interaction (*P*=1). Red bars indicate significant genotype differences (post-hoc independent two-tailed student *t*-test, *P*<0.05).

The spectral dynamics of the waking state in the 24h following Nestlet addition were first globally assessed in ECoG power heatmaps (Fig. 6b-c). This corroborated the existence of two nocturnal phases of heightened arousal, and, surprisingly, suggested that CKOs express a waking state higher in θ and fast-γ activity than do CTR littermates in early night (ZT12-15), when Nestlet novelty is maximal, and again in the last 2h of the night (ZT22-24), preceding sleep.

Because only the late-night phase was associated with sustained nestbuilding activity, we focused our spectral analysis on waking of the night’s last 2h (ZT22-24, Fig. 6d-e). To extract the electrocortical features specifically set in place in response to the nesting condition, we contrasted the ECoG power density values in waking during ZT22-24 of the nesting night to those in baseline waking in the same circadian time window. This confirmed that, conversely to EW and CC-induced waking, CKO mice’ wakefulness in late-night exhibit enhanced θ, as well as enhanced fast-γ (50-80 Hz) activity, compared to CTR mice (compare Fig. 6e, to 3f and 5d). Enhanced θ and fast-γ power in late-night furthermore coincides with a higher waking content (min/h) relative to CTR mice (Fig. 6a), that video data suggest may be related to nestbuilding activity.

Nestbuilding-associated waking of CKO mice exhibited moreover enhanced β (15-30 Hz) relative to CTR mice (Fig. 6e), as had been observed in CKO mice’ EW (Fig. 3f), and enhanced slow-γ (30.25-40.75 Hz) power (Fig. 6c,e). Elevated β and slow-γ synchrony are observed in dopamine-depleted rodents, and Parkinson disease (PD) patients in correlations with movement disorders^61,62^.

Because enhanced locomotion is part of the active waking response, it was also examined in this nesting paradigm. Surprisingly, CKO mice’ enhanced θ and fast-γ oscillatory responses were not matched by enhanced LMA. CKOs exhibited diminished LMA compared to CTRs, in early and late night (Fig. 6a), possibly reflecting stationary position assumed for Nestlet exploration/nestbuilding.

In summary, two paradigms of enhanced arousal under stress (EW and CC), and one ethologically-relevant, rewarding behavioral context (presence of nesting material prior to sleep), are found to elicit altered waking electrocortical responses in CKO mice relative to CTR mice, with, respectively, diminished, or enhanced θ and fast-γ activity. Figure 7 summarizes quantitatively these differential dynamics in waking θ and fast-γ oscillations. Because we here address the behavioral context-dependence of the dynamics of waking spectra, ECoG power is expressed relative to the average baseline (ZT0-24) waking ECoG activity in each frequency band. Nocturnal nesting-associated activity, and diurnal threat-avoidance (during ‘gentle handling’), or anxiety-related exploration (after CC^63^ appear therefore to induce highly divergent modes of HCRT-to-NA circuit processing and network oscillatory activity. Our data moreover suggest that these divergent oscillatory modes may alternate in the course of a day. CKO mice exhibit, as mentioned earlier, spontaneous periods of θ and fast-γ-enriched wakefulness in baseline dark phase, and, in a particularly pronounced manner, during the recovery night that follows EW (Fig. 4b).

**Figure 7.**
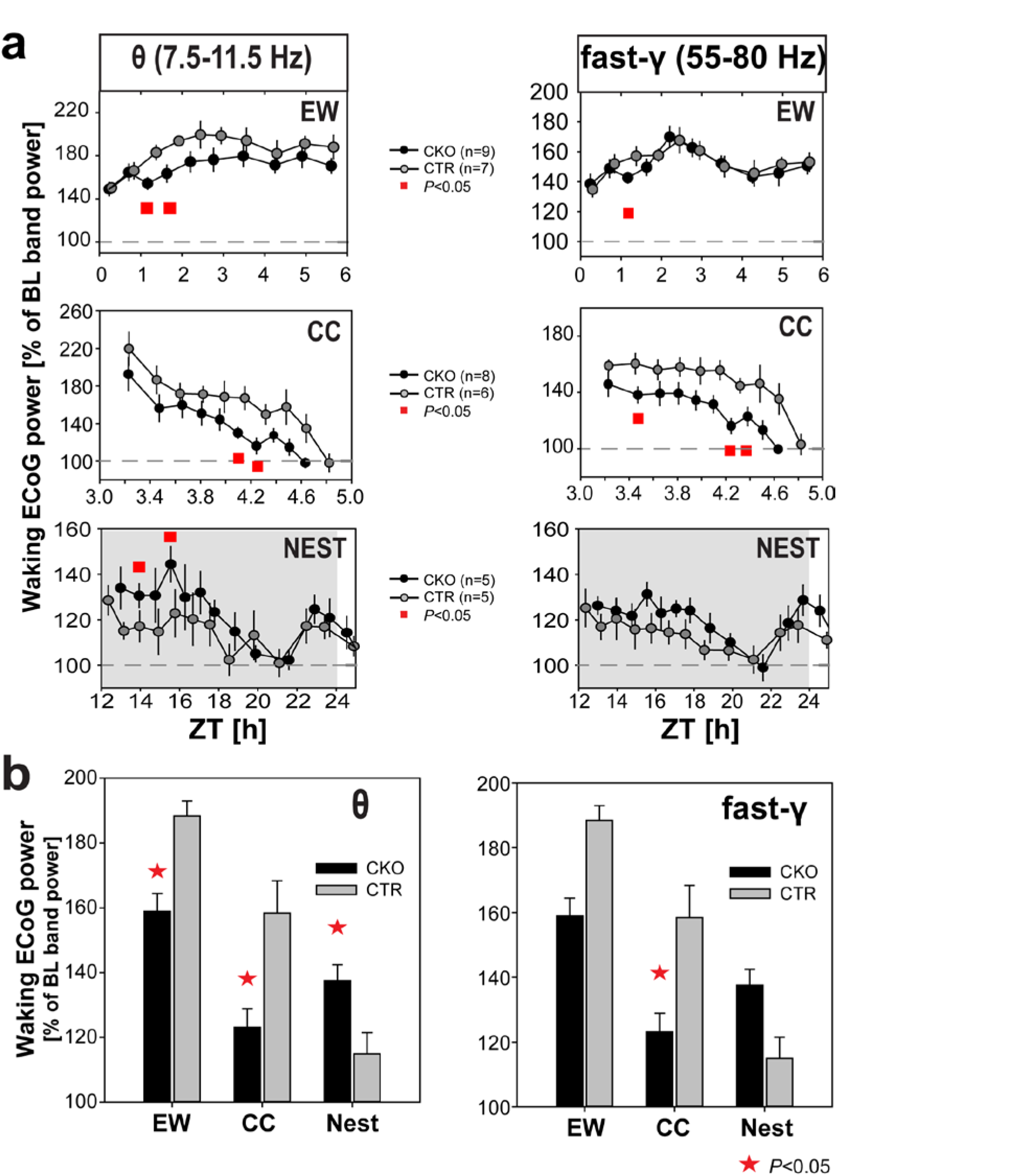
Bidirectional and context-dependent θ and fast-γ oscillatory responses of *Hcrtr1^Dbh-CKO^* (CKO) and *Hcrtr1^Dbh-CTR^* (CTR) mice in three paradigms of enhanced arousal, in adverse (EW, CC), or rewarding (NEST) environments. (**a**) Waking ECoG θ (7.5-11.5 Hz; *Left*), or fast-γ (55-80 Hz; *Right*) power is plotted across time in three behavioral paradigms: 6-h EW (ZT0-6; Top), from cage change (CC at ZT3) to sleep onset (*Middle*), or in the night following provision of a Nestlet at dark onset (ZT12; *Bottom*). Because we address the context-dependence of θ and fast-γ dynamics, ECoG power in each band is expressed as % of its average value in baseline waking (ZT0-24). For each time-interval, power values are calculated 0.25 Hz-frequency bin by 0.25 Hz-frequency bin (as in heatmaps of Figs 4–6), and then averaged across the respective frequency ranges. An equal number of artefact-free 4-s waking epochs contributed to each time-interval for each mouse. The following intervals are plotted: EW, 10 intervals spanning 6h; CC, 10 intervals spanning ~2h; Nest, 18 intervals spanning 24h (the first 13 are depicted). For each paradigm, two-way ANOVA for factors genotype and time revealed that θ power varies significantly with the animal’s genotype (EW and CC: *P*<0.0001; Nest: *P*=0.0025), and across time (*P*=0.0001). While all time-intervals show the same θ power trend in a particular paradigm, significant genotype difference is found for two intervals (□, *t*-test *P*<0.05). Two-way ANOVA also indicates a significant genotype effect on fast-γ power for CC (*P*<0.0001), and Nest (*P*<0.035), and a significant time effect on fast-γ power in all 3 conditions (*P*<0.0001). Note how the dynamics of θ and fast-γ oscillatory activities evolve in concert. (**b**) Waking θ and fast-γ power responses in the two time intervals marked by □ in (a) are plotted in the three behavioral paradigms as % of their respective mean power in baseline ZT0-24 waking. (Left) ECoG θ power varies significantly with genotype (*P*=0.0129), and condition (*P*<0.0001), and shows a significant genotype X condition interaction (*P*<0.0003, two-way ANOVA). CKO mice exhibit a significantly lower θ oscillatory response relative to CTR mice in wake during EW and after CC, but a significantly higher θ oscillatory response following Nestlet addition (✱ *t*-test *P*<0.05). (Right) ECoG fast-γ power tends to vary with genotype (*P*=0.0507), and varies significantly with condition (P<0.0001), and shows a significant genotype X condition interaction (*P*=0.0091, two-way ANOVA). t-test indicates that CKO mice display a significantly reduced fast-γ power response after CC (^✱^*P*<0.05).

### SWS following stress-associated arousal is selectively deficient in slow-δ frequencies in *Hcrtr1^Dbl-CKO^* mice

SWS quality critically depends on preceding wakefulness quality, in particular on time spent in θ– and fast-γ-rich wake (TDW)^53^. We previously showed that SWS δ power, a widely used marker of sleep homeostatic pressure, which is classically modeled as a function of prior sleep/wake history and increases with time spent awake^64,65^, fails to reflect prior time awake in *Hcrt^ko/ko^* mice in baseline. We found that this is linked to the fact that, while they spend a normal time awake, *Hcrt^ko/ko^* mice’ time in TDW state is profoundly diminished relative to WT littermates, due to a reduced ability to sustain TDW for a prolonged time^53^. We further found that SWS following spontaneous waking is not only reduced in δ power in *Hcrt^ko/ko^* mice, but also spectrally skewed, with a much more pronounced deficit in the slow-δ (δ1, 0.75-2.25 Hz) oscillatory component, than in the fast-δ (δ2, 2.50-4.25 Hz) range. The significance of distinct δ frequency ranges in SWS remains unclear, however a correlation between SWS δ1 component and prior active wake and related potentiation events was reported in other studies as well^66^.

Unlike *Hcrt^ko/ko^* mice, *Hcrtr1^Dbh-CKO^* mice spend a normal time in TDW in baseline. However their waking ECoG during EW and after CC is diminished in θ and fast-γ activity relative to controls. Thus we hypothesized that subsequent SWS may also be affected, and spectrally differ from controls. Hence we analyzed SWS of *CKO* and *CTR* mice in baseline and following the 3 behavioral paradigms they were exposed to. To best capture the entire range of the sleep homeostatic response before it dissipates^65^, we restricted our analysis on the first 20 min of SWS following: baseline light onset (Fig. 8a), EW (Fig. 8b), and CC (Fig. 8c), and differentially examined δ1 and δ2 activity. This revealed that CKO mice exhibit a severe δ1 power deficit in SWS following EW (Fig. 8b), and after CC (Fig. 8c), while δ2 power remains essentially normal. Baseline SWS (Fig. 8a) in contrast did not show a significance δ1 power deficit. CKO mice’ SWS after EW and the nesting night was moreover enriched in β and slow-γ activity (Fig. 8b and d).

**Figure 8.**
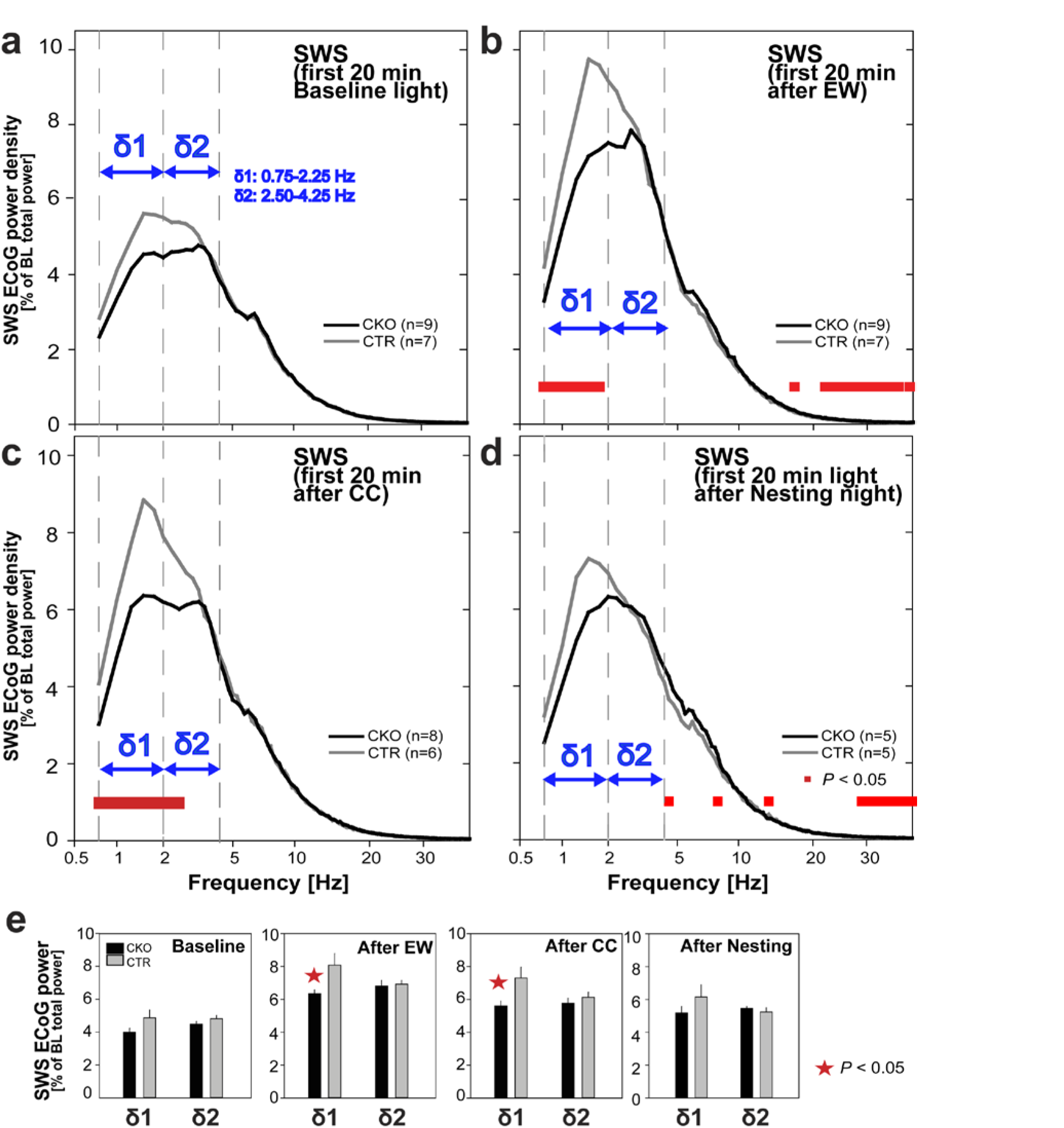
Slow-wave-sleep of *Hcrtr1^Dbh-CKO^* mice is deficient in slow-δ (δ1) oscillations following exposure to stressful stimuli. SWS ECoG spectra across 0.75-40.0 Hz are shown for the first 20 min of SWS (**a**) after light onset in baseline, (**b**) after EW, (**c**) after CC-induced waking, and (**d**) after light onset following the night in which nest material was provided. ECoG power is expressed as % of the same baseline reference as used for waking spectra of Figs 3b-c,e, 5c and 6d. Vertical guides delineate the two δ oscillatory subranges (δ1, 0.75-2.25 Hz; δ2, 2.50-4.25 Hz). SWS spectra of CKO mice differed significantly from those of CTR mice in all 4 conditions (a-d). Spectra depicted in a-c demonstrate significant genotype, and genotype X frequency effects across 0.75-40 Hz (two-way ANOVA, *P*<0.02 for all). SWS after nestbuilding (d) demonstrates a significant genotype X frequency interaction effect (two-way ANOVA, *P*=0.015), while factor genotype did not reach significance (P=0.82). In the δ range, only SWS following EW and CC (b-c) exhibited significant differences in CKO mice (post-hoc t-test, *P*<0.05), with in both cases a more profound deficit in δ1 power. No significant deficit in δ1 power in SWS following nestbuilding, or in baseline, were found. (**e**) Average SWS δ1 and δ2 activity in the 20 min intervals analyzed in (a-d) indicate that SWS δ1 activity is diminished only after EW and CC in CKO relative to CTR mice (^✱^, t-tests, *P*<0.05).

If CKO mice’ SWS δ1 deficit after stressful arousal is linked, in full or in part, to the θ and fast-γ power paucity of their preceding wake, through mechanisms similar to those operating in *Hcrt^ko/ko^* mice’ baseline SWS, CKO mice’ SWS after nestbuilding-associated waking, which conversely exhibits enhanced θ and fast-γ power, would not be expected to show an altered spectral distribution with underrepresented δ1 frequencies. CKO mice’ SWS following the nesting night was found to lack any δ, or selective δ1, power deficit (Fig. 8d), in accordance with our hypothesis.

Dynamic analysis of the two SWS δ oscillatory ranges across days 1-3 (Fig. S6c), and 8-9 (Fig. 6d) revealed δ1 to be diminished in CKO mice at all times, but more profoundly so following periods of maximal arousal (EW, and CC; Fig. S6c-dTop). Altogether, our data suggest that (1) HCRT-to-NA signaling is critical to mount normal θ and fast-γ responses to environmental stressors, (2) the content in θ and fast-γ frequencies of the waking ECoG affects not only subsequent SWS δ power, but also its spectral distribution, (3) waking enriched in θ and fast-γ frequencies induces preferentially a low-frequency component of δ oscillations in subsequent SWS.

## Discussion

The main finding of this study is that chronic inactivation of HCRT-to-NA signaling affects θ and fast-γ frequencies during waking, and their repercussions on subsequent slow-δ frequencies during SWS. θ and fast-γ oscillations are electrocortical markers of active wakefulness and motivated behaviors. They are thought to provide a temporal framework on which the timing of neuronal spikes can be measured and computed during the execution of complex behaviors such as navigation^67^. Induction of these two markers of arousal were diminished in CKO mice relative to control littermates during EW and following CC, whereas the opposite was observed during nesting-associated wakefulness preceding sleep.

### Scope of the *Hcrtr1^Dbh-CKO^* mouse model

In CKO mice, NA neurons lack HCRTR1 in their soma, dendrites, as well as synaptic boutons. HCRT induces depolarization and increased firing rate of NA^LC^ cells^16,18,20^, suggesting that HCRTR1 is present on NA^LC^ cell bodies or proximal dendrites, in agreement with the intense *Hcrtr1 in situ* signal over the LC^26^. The present study confirms at a cellular resolution that the *Hcrtr1* gene promoter is active in NA^LC^ neurons, as intense GFP-immunofluorescence was present in 98% of LC TH^+^ cells of CKO, but none of their CTR littermates. HCRTR1 can enhance neurotransmitter release through both spiking regulation and presynaptic effects^41–46,68^, and CKO mice are expected to be deficient in HCRTR1 expression and signaling not only in the NA nuclei, but also in their target tissues, such as the hippocampus, PFC, amygdala, ventral tegmental area (VTA), and the olfactory bulb^10,17^. However, the CRE-mediated genomic recombination of the *Hcrtr1*^flox^ allele activates expression of a cytoplasmic GFP in place of HCRTR1. Therefore, GFP distribution informs about cell identity, but not receptor’s intracellular distribution, and whether HCRTR1 is expressed in NA cell terminals remains uncertain.

The downstream effectors of HCRTR1 are known to be multiple and context-dependent. HCRTR1 can act through a great variety of signaling cascades, to mediate short-term changes in ionic conductances, or long-term changes in neuronal circuitry^24^. As HCRT enhances NA^LC^ cell firing, we interpret CKO mice’ phenotype at first approximation as the result of widespread loss in HCRT-induced NA signalling. HCRTR1 deficiency may however alter many other functions of NA neurons, such as signalling through other receptors, release of other neurotransmitters^69^, synaptic plasticity^70^, or gene expression, all of which contributing to the global phenotype. Further electrophysiological and molecular analyses of our mice are needed to evaluate these effects.

### Slowing of the oscillatory profile of wakefulness

Total time spent awake was normal in CKO mice in baseline conditions. Thus, although awakening induced by optogenetic stimulation of HCRT neurons was shown to be blocked by the concomitant photoinhibition of NA^LC^ cells^29^, loss of HCRTR1 in NA cells of CKO mice did not diminish total baseline waking, bout number or duration. This discrepancy may have several causes. First, whereas optogenetic manipulations acutely modulate neuronal activity, NA cells of CKO mice are thought to lose *Hcrtr1* gene function at the time the *Dbh-Cre* transgene is first expressed, i.e. in mid-gestation^36^. Compensatory mechanisms set in place during development may partially rescue CKO mice’ wake by upregulating other arousal pathways. For instance, increases in histamine cell number were reported in both narcoleptic patients and mouse models^71^. To examine whether NA^LC^ cell-specific *Hcrtr1* gene inactivation at an adult stage causes a more severe phenotype than the one we report in CKO mice, a *Th*-driven *Cre*-expressing viral vector may be injected in the LC of adult *Hcrtr1^flox/flox^* mice. Second, although under optogenetic settings the HCRT-to-NA^LC^ circuit critically mediates awakening, under physiological conditions, other HCRT targets may also elicit state transitions, and the HCRT-to-NA^LC^ circuit may mostly be critical under specific alerting contexts that are not met in our baseline conditions, or not at a rate sufficiently high to alter total time awake. Next, in the above optogenetic study, and upon HCRT infusion in LC *in vivo*^20^, NA^LC^ cell activation and state transition may not only be caused by released HCRT binding HCRTR1 on NA cells, but also HCRTR1 on synaptic terminals of a third LC cell type, e.g. glutamatergic neurons^72^, thus enhancing glutamate release, NA cell firing, and thereby the probability of awakening. The latter mechanism would be preserved in CKO mice.

While CKO mice’ waking time was not changed, its spectral quality was. Baseline waking was enriched in δ (1.25-2.75 Hz) frequencies, and upon EW and CC, in δ and inter-δ/θ frequencies, both of which reflect homeostatic sleep pressure in rodents and humans. Waking δ activity moreover correlates with declining cognitive performance during prolonged waking, and elevation of δ, θ/α, and β power, while γ power fades, correlate with subjective sleepiness^73–79 55 80^. Thus an increase in slow (infra-θ) oscillations in CKO mice suggests a less alert waking state, which appears to affect waking globally, as active waking (TDW) is also excessively δ-rich. Mechanistically, waking δ activity may stem from localized SWS network activity occurring in the brain of a globally awake, but sleepy, animal^81^.

### Increased waking β and slow-γ activity

In times of enhanced arousal, such as during EW and nestbuilding, CKO mice display enriched β, or β and slow-γ (~15-45 Hz) power. Dopamine-depleted rodents also exhibit increased β/slow-γ activity while performing particular tasks^62,82^. Whether alterations in dopaminergic transmission are also present in our mice, potentially resulting from diminished dopamine release from NA^LC^ cells^69^, or from downregulation of a HCRT-NA^LC^-DA^VTA^ circuit, deserves further investigation. Its association with impaired motor control also merits further investigation. Pathological β oscillations are present in Parkinson’s patients, as an ‘antikinetic β band’ correlated with movement disorder^61^. *Hcrt*-KO mice also show enhanced β/slow-γ activity in the minute that precedes cataplexy, which is characterized by indices of both electrocortical and behavioral arousal, with vigorous activity, including nestbuilding^83^. Dysregulated motor events can also directly precede cataplexy, supporting the hypothesis that our mice’ β/slow-γ-enriched wake is related to PD’ antikinetic rhythm. Furthermore, the increase of β/slow-γ frequencies associated with diminished θ we observe in periods of intense arousal aligns with observations in rats, in which LC activation caused a suppression of β/slow-γ (12-40 Hz) frequencies, while enhancing θ oscillations^84^. LC activation was suggested to mediate this effect by repressing inhibitory hippocampal interneurons, thus enhancing conditions for synaptic potentiation and learning.

### Diminished induction of θ and fast-γ frequencies under stress

In stressful conditions, as experienced by the mice during sleep deprivation and following CC, CKO mice had a reduced ability to mount a normal arousal response, consisting of a sharp θ spectral peak associated with a fast-γ power surge. Numerous types of stress are associated with activation of HCRT neurons^8^, and we previously showed that HCRT is essential to stabilize θ– and fast-γ (θ/fast-γ) -enriched wakefulness^53^. Emergence of a narrow-bandwidth θ rhythm may involve a process enhancing or stabilizing θ oscillation synchronization in hippocampal or neocortical networks. NA^LC^ activation indeed enhances θ power and resets θ synchronization in hippocampus and neocortex^85–87^. It was recently proposed that LC neuron spiking, NA release, and θ oscillations, are temporally and mechanistically linked in the process through which the LC resets network activity, and mediates attentional shift, in response to salient stimuli^88^. Our data add to this view the role of the stimulatory input contributed by the HCRT-to-NA^LC^ pathway in mediating LC activation, in particular in stress-associated contexts. Furthermore our results are consistent with the reports evidencing HCRT involvement in θ wave generation or maintenance *in vitro*^89,90^, and *in vivo*^91,92^. HCRT also regulates θ–associated plasticity in the hippocampus by a presynaptic mechanism involving NA signaling^93^. From a cellular standpoint, HCRT stimulates the septal parvalbumin-expressing (PV) GABAergic neurons. These engage into the septo-hippocampal GABAergic networks that are thought to pace hippocampal principal cell synchrony, thus generating the θ-rhythm^94,95^. Our findings confirm and extend the evidence linking HCRT, NA and hippocampal θ oscillations, warranting behavioral investigation of hippocampal NA-dependent forms of learning in CKO mice.

Although we do not have θ-γ phase coupling data, a salient finding of our study is the consistent association between θ and fast-γ dynamics. In fact, the hippocampal networks involving PV neurons modulated by NA are also implicated in coupling θ rhythm generation with γ (35-85 Hz) waves^96–98^, and PV cells may provide a mechanism for NA^LC^ support of phase-coupled rhythms^99^, and its regulation by a HCRT input. In addition, NA is known to increase γ (30-60 Hz), and decrease δ activity when injected in the basal forebrain (BF) of rats. The effect is mediated by NA depolarizing activity on BF cholinergic neurons^100^. These cells project cortically and their implication in stress-induced, HCRT-dependent, arousal merits further investigation.

### Inverted θ and fast-γ responses in nesting-associated vs. stress-associated wakefulness

The ECoG exhibits a sharp spectral peak at θ frequencies (8-10 Hz), and a concomitant rise in fast-γ power when mice engage in intensive voluntary behaviors, both in threat-avoiding, or reward-seeking conditions. We exposed CKO and CTR mice to both types of contexts. Gentle handling during EW, and CC both induce a threat-avoiding-like response^63^, while nest material induces vigorous nestbuilding, and is rewarding^59^. CKO mice exhibited abnormal electrocortical responses in both types of conditions, albeit in opposite directions. They showed lower θ and fast-γ activity than controls in enforced and CC-induced waking, but θ and fast-γ power was higher than controls in late-night nestbuilding-associated waking preceding sleep. What could explain these opposite responses?

While arousal induced by both aversive and appetitive stimuli engage high θ and fast-γ oscillatory activity, they likely use distinct neurocircuits. Both are however expected to be modulated by HCRTR1 signaling^8^. Our prior work has demonstrated that θ/fast-γ-enriched waking is differently regulated when waking is spontaneous, or enforced, and differently affected by HCRT deficiency^53^. Considering the high complexity of HCRTR1 neurotransmission at both the circuit and cellular levels, a multitude of factors may contribute to context-dependent modulation of θ and fast-γ oscillations. As a working hypothesis, we focus on fear-(amygdala), and reward-(VTA) related structures, which receive both direct projections from HCRT and NA cell groups^101–103^.

A serial HCRT-NA^LC^-lateral amygdala (LA) circuit in which intra-LC HCRTR1 signaling features a key role was recently shown to be critical for formation, maintenance, and extinction of fear memory^104–106^. Blocking LC HCRTR1 activity reduced, while upregulating HCRT signaling within the LC enhanced, freezing behavior, a readout of fear memory. Furthermore, inhibition of HCRTR1 signaling within the amygdala facilitated fear memory extinction^107^.

Therefore, loss of HCRTR1 at the level of NA^LC^ cell bodies, and/or at synaptic terminals in the amygdala, may contribute in blunting the fear response of CKO mice. It may be relevant to note in this context that reduced amygdalar activity during aversive conditioning was reported in human narcolepsy^108^. CKO mice’ deficient θ and fast-γ oscillatory response upon EW and CC may indicate a weakened alerting response to threat. This is consistent with the observation that CKO mice’ waking ECoG spectra during EW and after CC are markedly enriched in frequencies associated with diminished alertness and sleepiness (δ, inter-δ/θ and β/slow-γ), compared to CTRs. A reduced fear response may contribute, conversely, to CKO mice’ enhanced behavioral activation during the self-motivated activities mice spontaneously engage in nocturnally, or when nest material is provided, causing the increased waking time, and θ/fast-γ oscillatory response they express in late-night phase, which is rich in pre-sleep nestbuilding activity. Assessment of CKO mice’ fear response will however require more elaborate behavioral and physiological measures of fear.

Tests for impulsivity and risk-taking behavior may furthermore complement assessment of CKO mice fear response. Increased impulsivity, and patterns of disadvantageous decision making were reported in narcolepsy^109,110^. These traits were hypothesized to result from emotional blunting associated with HCRT deficiency. Thus, although increased sleepiness is a cardinal symptom of narcolepsy, other symptoms such as increased impulsivity, or binge-eating behavior, might suggest forms of hyperarousal^110,111^.

At the cellular, mechanistic level, the bidirectional θ/fast-γ response of our CKO mice may involve multiple NA cell signaling pathways. Again we focus on one example. In addition to promoting NA release from axonal terminals in target tissues, HCRT is also reported to trigger and potentiate NA secretion non-synaptically, from the somata and dendrites of NA^LC^ cells^112,113^, in a process that is HCRTR1-, and NMDA receptor-dependent^112^. Therefore HCRTR1 may not only modulate NA input to targets, but also proximal regulation of LC neuronal activity. Because binding of locally secreted NA to NA^LC^ cell autoreceptors represses NA^LC^ activity^114^, somatodendritic secretion provides a negative feedback regulatory pathway by which HCRTR1 may inhibit LC activity. NA somatodendritic secretion was indeed suggested as a potential therapeutic target to depress the NA^LC^ hyperactivity that results from opioid withdrawal^112,115^. Because somatodendritic NA secretion has distinct properties compared to classical synaptic transmission, including delayed kinetics and a tight dependence on the frequency of the incoming stimulation^116^, it may be favored over the latter in some behavioral contexts, possibly contributing to the enhanced behavioral activation and θ/fast-γ response of CKO mice in nocturnal nesting conditions.

### Chronic loss of HCRT-to-NA signaling does not cause overt cataplexy

HCRT deficiency is known to cause the life-long disease narcolepsy with cataplexy. Ample evidence in canine models implicated the NA system in the control of cataplexy, leading to the NA hypoactivity hypothesis of cataplexy^117^. In narcoleptic dogs, NA cells cease firing at the onset of cataplexy episodes and resume spiking at cataplexy offset^118^, suggesting NA^LC^ activity is critical to couple arousal with muscle tone. Burgess et al.^119^ suggested that cataplexy is caused by loss of HCRT control of the NA system, which is dispensable for consciousness, but needed for the associated muscle tone. Although a comprehensive video-based behavioral assessment of CKO mice is yet to be performed, these animals did not demonstrate obvious electrocortical, or behavioral evidence of cataplexy^83^, in concordance with more recent studies^32,120^.

### HCRT-to-NA signaling deficiency affects the spectral quality of SWS

We found that SWS following stressful wake is selectively deficient in slow-δ (δ1) power in CKO mice. This is reminiscent of our observation of a similar δ1 deficit in the SWS of *Hcrt^ko/ko^* mice following spontaneous waking. The latter was associated with a profound deficit of *Hcrt^ko/ko^* mice in expressing θ and fast-γ activity during baseline waking, due to a profoundly reduced ability to sustain spontaneously electrocortical arousal (the TDW state), as reflected by a much lower average TDW bout duration compared to controls^53^. Here, we show that CKO mice express a waking state globally reduced in θ/fast-γ activity under stressful conditions, although TDW episode duration is essentially normal (Fig. S3). Thus a deficit in δ1 activity is selectively observed in SWS subsequent to θ/fast-γ-power-impoverished waking. We have shown that time spent in θ/fast-γ-rich/TDW state was the key driver of the sleep homeostatic response, since its proxy SWS δ power was seen to correlate with prior time spent in TDW, rather than in all waking^53^.

A dependency of SWS δ1 on prior waking θ/fast-γ activity is strengthened by our observation that CKO mice’ SWS after nesting-associated wake, which, inversely to EW and CC waking, exhibit enhanced θ/fast-γ activity relative to controls, shows normal δ1 power. Our findings are also in line with the report that recovery SWS of rats that were depleted from cortical NA show a similar selective δ1 deficit^66^. Moreover, cortical expression of a set of synaptic activity-related genes was found to be diminished in these rats during EW^66^, as we also found to be the case in *Hcrt^ko/ko^* mice in baseline dark phase^53^.

Altogether these studies suggest that the δ1 component of SWS activity has a privileged homeostatic link with, and dependency upon, θ/fast-γ oscillatory activity in prior waking. θ/fast-γ-dependent potentiating events, or other plasticity changes within cortical circuits which get facilitated by HCRTR1-mediated noradrenergic signaling during waking, may be causally linked to δ1 oscillations in subsequent SWS^121^. Our study suggests that HCRTR1-dependent NA signaling contributes to the homeostatic weight of waking, in particular as it is reflected in δ1 power in ensuing SWS. This hypothetically predicts that SWS δ1 oscillations mediate a specific, yet unknown, function in SWS-associated recovery processes.

The relation between the slow-δ (δ1) component of SWS and the cortical slow (<1 Hz) oscillation of alternating UP and DOWN states in cortical neurons^122^, is unclear. Whereas the slow oscillation was originally identified through intracellular recordings, δ1 is a Fourier–derived rhythmic component of the ECoG. Two features of δ1 nevertheless suggest that δ1 and the slow (<1 Hz) oscillation are distinct: (i) the latter is reported to peak at ~0.3 Hz, whereas our δ1 rhythm is distinctly faster (0.75-2.25 Hz); (ii) while we clearly document the wake/sleep-dependent homeostatic regulation of δ1, whether the slow oscillation is under homeostatic regulation is controversial. Both early and recent EEG studies in humans have suggested that the oscillatory component below 1 Hz is unresponding to sleep deprivation^123,124^, strengthening the hypothesis that the two rhythms have different origins. This important question warrants further studies.

### PS state consolidation

Lastly, while CKO mice’ episode number, duration, and total time spent in waking or in SWS were normal in the conditions we tested, the mice exhibited a modest but significant increase in PS stability, resulting in a shift towards longer PS bouts. Both HCRT and NA are known to have PS-antagonizing activities. Inhibiting NA signaling enhances, while increasing NA represses, PS^125^. Narcoleptic patients show an increased propensity to express the PS state^126^, and the same is true for Hcrt deficient mice in some conditions^53 83,127^. The direct administration of HCRT-1 peptide into the LC of rats was shown to cause a decrease in both PS episode number and duration^20^. In our mice, chronic NA cell HCRTR1 deficiency appeared to increase PS maintenance, but PS initiation was unchanged.

### Conclusions

Brain processes that alter the generation or the stability of θ and fast-γ waves appear particularly active in presence of ethologically-relevant stimuli, of either repulsive or attractive valence, of importance for survival. These dynamics can result from neuromodulation of circuits by wake-promoting molecules, such as NA and HCRT, and their receptors. Many of the mechanisms, both at cell and circuit level through which these molecules act remain elusive. While HCRT is essential for state stability^128^, including stability of the θ/fast-γ-enriched TDW state^53^, we suggest that the HCRT-to-NA circuit is a context-dependent modulator of the different spectral domains in which brain networks can engage, and that differentially assist and organize brain operations in wake, and their processing in sleep.

The understanding of how HCRT neurotransmission affects noradrenergic function has clinical relevance beyond implications for narcolepsy and cataplexy, as HCRTR antagonists are being developed as novel therapeutic agents not only for insomnia, but also in several neuropsychiatric disorders, such as anxiety, depression, and various forms of hyperarousal, including cocaine-withdrawal-, and post-traumatic-stress-related hyperarousal^129^. Furthermore, HCRTR1 ability to modulate LC neuronal activity is providing a novel therapeutically relevant tool in several neurodegenerative disorders in which LC neuronal damage appears to be an early event^130^. Many remaining challenges exist for HCRTR antagonists’ use in clinics, in particular because of the pleiotropy of HCRT functions, affecting not only arousal, but also e.g. food uptake, motivation and mood. The bidirectional effects of *Hcrtr1* disruption in the NA system we report warrant further studies also for these reasons. Thus our mouse model, and others that similarly inactivate a specific neuromodulator receptor in selective cell populations, will contribute in dissecting the mechanisms by which these molecules shape brain states and behavior.

## Methods

### *Hcrtr1* gene engineering: creation of a CRE-dependent conditional KO (*Hcrtr1^tm1.1Ava^*) allele

#### Targeting strategy

We inserted a 5’loxP site in the 5’ untranslated region of the first coding exon (exon 3 according to^131^) of *Hcrtr1*, 27 bp downstream of exon 3’s splice acceptor site, and 12 bp upstream of the initiation codon. The 3’ loxP site was inserted midway in intron 4 as a *loxP-Kozak-Gfp-rabbit globin p(A)-FRT-Neo-FRT* cassette. The inter-loxP distance is 1.1 kb.

#### Mutagenesis

The targeting vector for homologous recombination (Fig. 1a) was built using plasmid artificial chromosome (PAC) *RP23-12J1* (BACPAC Resources Center, Oakland, CA, USA) a PAC from a *C57BL6/J* genomic DNA library containing *Hcrtr1*. Homologous arms spanning in total 10.0 kb were synthesized by PCR amplification of PAC *RP23-12J1* using LA Taq (Takara), and Accuprime High-Fidelity (invitrogen) DNA polymerases. The targeting vector was linearized and electroporated in IC1, an embryonic stem (ES) cell line derived from *C57BL/6NTac* mice (ingenious targeting laboratory, Ronkonkoma, NY, USA). Single colonies were screened by Southern blot analysis using probes external to the targeting vector in both 5’ and 3’, and a *Gfp* internal probe. Two ES cell clones having undergone correct recombination events were identified and injected into *BALB/cAnNHEW* blastocysts. Resulting chimeras were mated to *Tg(ACTFLPe)9205Dym* transgenic mice^132^ to excise the FRT-neo-FRT cassette. The neo-excised final conditional KO allele, *Hcrtr1^tm1.1Ava^* (MGI: 5637400), is referred hereafter as *Hcrtr1flox (http://www.informatics.jax.org/allele/MGI:5637400*).

### Generation of noradrenergic cell-specific *Hcrtrl*-KO (*Hcrtr1^Dbh-CKO^*) mice, and evidence of tissue-specific, C redependent *Hcrtr1^flox^* allele recombination and genomic deletion

To inactivate *Hcrtr1* in NA cells, transgenic *Tg(Dbh-icre)^1Gsc^* mice, which carry a PAC transgene with *Dopamine-β-hydroxylase gene* (*Dbh*) sequences driving Cre^36^, were mated with *Hcrtr1^flox^* mice. Two PCR assays flanking the loxP sites of *Hcrtr1^flox^* were designed to amplify a diagnostic fragment of the recombined allele, and a 1.1 kb larger fragment (inter-loxP distance) from the unrecombined allele. Genomic DNA was prepared from tissue punches made in brain slices containing the LC, or other brain areas. Genomic DNA from the LC area of *Hcrtr1^Dbh-CKO^* mice amplified the diagnostic recombined fragments, but not from the LC of *Hcrtr1^Dbh-CTR^* littermates, neither from neocortex, or ear tissue of either CKO or CTR mice. CKO mice’ PCR products were gel purified and sequenced in both orientations, confirming accurate recombination at the nucleotide level in 2 *Hcrtr1^Dbh-CKO^* animals.

### Generation of Hcrtr1 KO/GFP-reporter (*Hcrtr1^tm1.2Ava^*) mice

*Hcrtr1^flox/flox^* mice were mated to *Tg(EIIa-cre)^C5379Lmgd^* transgenic mice^49^, which express Cre at embryonic preimplantation stages. Offspring that stably transmitted the recombined (*Hcrtr1^tm1.2Ava^*) allele were selected for further breeding, and *Tg(EIIa-cre*) was segregated out. Genomic DNA extracted from two animals was sequenced to confirm accurate CRE/loxP recombination at *Hcrtr1*. Resulting mice carry the recombined KO/GFP-reporter (*Hcrtr1^tml.2Ava^* MGI: 5637401, or *Hcrtr1^KO-Gfp^*) allele in all cells (*http://www.informatics.jax.org/allele/MGI:5637401*). Because the GFP coding sequence in this allele does not, unlike the HCRTR1 coding sequence that it replaces, carry a N-terminal signal peptide, it generates cytoplasmic GFP, which distributes through soma and neurites. Thus GFP expression reflects *Hcrtr1* promoter expression, but does not inform about HCRTR1 protein intracellular distribution.

### Experimental mice

All mice derived from mating between *Hcrtr1^flox/flox^* mice, with one of the two parents hemizygous for *Tg(Dbh-Cre*). These crosses generate two offspring groups: *Hcrtr1^flox/flox^; Tg(Dbh-Cre*)^+^ mice, in which NA cells express Cre (*Hcrtr1^Dbh-CKO^*, or CKO mice), and *Hcrtr1^flox/flox^* littermates, which do not express Cre (*Hcrtr1^Dbh-CTR^*, or CTR mice). All analyses relied on pair-wise phenotype comparison between these two groups. All mice share a *C57BL/6NTac* X *C57BL/6J* mixed genetic background. All ECoG data are derived from 10 to 13-week-old males (weight 27-31 g). Mice were individually housed with food and water ad libitum under an LD12:12 cycle (lights-on, i.e., Zeitgeber Time ZT0, at 08:00 AM). All experiments were performed in accordance with the Swiss federal law, and using protocols approved by the State of Vaud Veterinary Office.

### ECoG/EMG, video and locomotor activity (LMA) recording

Electrodes for differential fronto-parietal ECoG and EMG recording were implanted as described^133^. Mice were allowed to habituate to the EEG-setup for a minimal of 10 days, before the recordings were started (Fig. 2). Hardware (EMBLA A-10) and software (Somnologica-3) were from Medcare Flaga (EMBLA, Thornton, USA). ECoG and EMG signals were amplified, filtered, analog-to-digital converted, and stored at 200 Hz. Behavioral states (wake [W], slow-wave-sleep [SWS], and paradoxical sleep [PS]) were scored visually in 4-second (4-s) epochs using published criteria^133^. Theta-dominated-waking (TDW) epochs were identified among waking epochs using an ECoG-based algorithm as described^53^. Behavioral monitoring was done using infrared video-cameras^83^. Locomotor activity was assessed using top-mounted passive infrared motion detectors (Visonic Ltd, Tel Aviv, Israel), and analyzed at 1-min resolution using ClockLab software (ActiMetrics, IL, USA).

### ECoG power spectral density (PSD) analysis

The ECoG signal was subjected to discrete Fourier transform to yield power density spectra (0.75–90 Hz) in 4s epochs (0.25 Hz frequency resolution; Hamming window function). For each behavioral state and mouse, a mean ECoG spectrum was obtained by averaging spectra of all artefact-free epochs of that state and flanked by same-state epochs^134^. To account for individual differences in absolute ECoG power, power density is expressed as % of a baseline ECoG power reference (‘% of BL total power’), which is calculated for each mouse across two days, all states, and summation of the 0.75-40 Hz frequency bins. This reference is weighted so the relative contribution of each state is identical for all mice^134^. Reference values for CKO and CTR mice did not differ (CKO vs. CTR 1181± 127 uV2 vs. 1181± 120 uV2, t 14= 0.00105, P=0.99). Power density values across 47.5-52.5 Hz were excluded due to the 50 Hz AC power line noise. Power density values across 71.25-77.25 Hz were also excluded due to artefacts of unknown origin present in some mice (EW: 6/16, 3CKO:3CTR; CC: 4/14, 1CKO:3CTR; Nest experiment: 4/10, 1CKO:3CTR). To rule out that some of the power spectral density changes observed in CKO relative to CTR mice are an effect of referencing power density values to the individual’s mean ‘total power’ as described above, we also compared the spectral density profiles of the two genotypes using their absolute ECoG power values. This yielded significant spectral differences between CKO and CTRs in the same frequency bands as the ones obtained with the ‘total power’ reference. Greek symbols were used to define specific ECoG frequency ranges as follows: δ (1-4 Hz), inter-δ/θ (4-7 Hz), θ (7.5-11.5 Hz), β (15-30 Hz) slow-γ (γ-1: 32-45 Hz), and fast-γ (γ-2: 55-80 Hz). In Fig. 3e inset and Supplementary Table S2, the dominant frequency of θ oscillations (θ peak frequency, TPF) was evaluated in each mouse as the frequency at which ECoG power density in the average waking spectral profile was highest in the 5-10 Hz range^135^.

### Temporal dynamics of integral PSD in specific frequency bands

The dynamics of ECoG activity within specific frequency ranges in waking across various contexts and periods of time were analyzed by dividing the period concerned into a same number of time intervals (‘percentiles’) for all animals, each percentile contributing an equal number of 4s epochs scored as waking. The number of time intervals, and thus of data-points, was adjusted according to the prevalence of wakefulness (e.g., 6 in baseline light phase, 12 in dark phase, 8 during the 6-h EW, and 4 in recovery light phase, for Fig. 3d). For each time interval, the mid-points were averaged among animals of a given genotype, and mapped back onto real time. Because time-spent-awake (e.g. from cage change to SWS-onset) may differ between genotypes, data-points of each genotype do not necessarily coincide in time (x-value). ECoG power density in each frequency band was expressed relative to its average value in baseline wake (ZT0-24; Figs 5g, 7a), or relative to the ‘baseline total power reference’ described above (Fig. 3d). Genotype differences in ECoG power dynamics were statistically evaluated according to the percentile sequence, i.e. CKO and CTR mice’ ECoG power values were compared in each successive time percentile.

### 3D (ECoG Power:Frequency:Time) heatmap analysis of wakefulness

To analyze the time dynamics of the entire ECoG spectrum (0.75-90 Hz) in waking, power density in each 0.25 Hz frequency bin was first expressed relative to its average level in baseline light phase waking across the ZT8-12 time interval, then averaged across each time interval, and individual of each genotype. Results are visualized as a 3D ‘heatmap’, with larger changes in power density appearing as warmer colors. It is important to note that, in heatmaps, (as in all analyses depicting ratios between ECoG power values), the frequencies at which the ECoG power value peaks (warm-colored areas) do not represent the precise frequencies of the brain’s actual most powerful oscillations, but rather the frequencies at which power densities maximally differ with reference baseline values. These frequencies are slightly right-shifted (higher), relative to the frequencies of the actual brain oscillatory rhythms that dynamically respond to the environmental contingencies under analysis. This principle is illustrated in Fig. S4.

### Behavioral exposure

Cohorts of CKO and CTR (n=9:7, CKO:CTR) littermate mice were sequentially exposed to four experimental contexts, as their ECoG/EMG data were acquired: (i) baseline (undisturbed) conditions, (ii) EW, (iii) nestbuilding, (iv) transfer to fresh cage. Only uninterrupted high-quality recordings were considered for analysis. Hence, (i-ii): n=9:7; iii: n=5:5; (iv): n=8:6 (CKO:CTR) animals were analyzed.

#### Spontaneous and enforced wakefulness

Mice were recorded for 3 continuous days, consisting of two undisturbed (baseline) days (BL1+BL2), a 6-h period of ‘gentle handling’^133^ initiated at light onset (ZT0) of the 3rd day to maintain animals awake, and a 18-h ‘recovery’ (undisturbed) period. To avoid sleep during EW, tissue paper was introduced in the cage at ZT2 and ZT5, and midway through EW (ZT3), mice were transferred into a fresh cage.

#### Nest material interaction

Following the day of the EW, mice were left undisturbed for ≥2 days. At dark onset (ZT12) of day 6, a square of highly packed shreddable cotton (NestletTM, Ancare, Bellmore, NY, Ref. Nr. 14010) was introduced in the cage. Mice were then left undisturbed through the night until light onset and assessment of nest morphology. All nests (CKO n=5; CTR n=5) were found to score at least 3 according to Gaskill et al.59’s nestbuilding scale.

#### Cage change (CC)

Following nest assessment, mice were left undisturbed for another ≥2 days. On ZT3 of day 9, mice were transferred from their homecage where the nest had been built to a fresh cage. Latency to SWS-onset was defined as the time until the first SWS episode lasting ≥2 min and interrupted by ≤3 epochs scored as wakefulness.

### Immunofluorescence

~16-wk old mice were deeply anesthetized with sodium pentobarbital (100 mg/kg, i.p), and transcardially perfused with 4% paraformaldehyde (pH 7.4). Brains were quickly removed, post-fixed in the same fixative for 2h at 4°C, then immersed successively in 15% (1h) and 30% sucrose (o/n) at 4°C, frozen and stored at −80°C until use. Coronal sections (20-μm) were collected on SuperFrost-Plus glass slides, blocked in 2% BSA, 5% normal donkey serum, 0.3% TritonX-100 in TBS (pH 7.5) for 30 min at r.t., and antibodies were applied in 1% BSA, 0.2% TritonX-100 in TBS, and incubated o/n at 4□C. When double staining involved the HCRTR1 antibody, an additional ‘antigen-retrieval’ step in Sodium Citrate buffer (pH 6.0) buffer at 55oC for 7 min was applied before blocking and primary antibody incubation. Antibodies were: anti-tyrosine hydroxylase (TH) from mouse (Incstar, Cat#22941; 1:5000), anti-green fluorescent protein (GFP) from chicken (Aves Labs, Cat#1020; 1:500), anti-CRE from rabbit (Novagen, Cat#69050-3; 1:500), and anti-HCRTR1 from rabbit (Origene Cat#TA328918; 1:100-1:500). This HCRTR1 antibody was raised against a 15-aa peptide from the third intracellular loop of rat HCRTR1, i.e. a region downstream of the 126-aa deletion mediated by CRE/loxP recombination of the floxed *Hcrtr1* allele Secondary antibodies were donkey IgGs coupled to Alexa-594, or -488 fluorophores, and were incubated at a 1:500 dilution for 1h at r.t.

### Confocal microscopy

Images were acquired on an inverted Zeiss LSM710 confocal laser-scanning microscope (405, 488, and 561 nm lasers) using a 40x oil objective (EC plan-Neofluar 40x/1.30 Oil DIC M 27). For each animal used for cell quantification, sections collected at 4 Bregma levels within the LC (representing approx. 1 out of 9 sections) were used (−5.34, - 5.40, −5.52, −5.68 mm from Bregma^136^. Confocal images covering the entire LC TH+ cell field were acquired at 8-bit image depth and a frame of 1,024 × 1,024 pixels, and tiled together using ZEN software. A square frame centered within the LC field was drawn (1,024 × 1,024 pixels at 40×, as shown in Fig. 1b-f), and immunoreactive cell counts were evaluated within that frame using ImageJ software. Images were minimally processed in the same manner for the two genotypes.

### Statistics

Data were assessed for normal distribution using Shapiro-Wilk normality test, and nonparametric Mann-Whitney statistics were used if the test failed. Otherwise, timecourse and ECoG power spectral density analyses were assessed first by two-way ANOVA for factors ‘genotype’ and ‘time-of-day’, or ‘genotype’ and ‘frequency bin’, and their interaction. Only if a significant effect was found for the factor genotype, or/and the two-factor interaction, were genotypes contrasted by independent t-tests. Within-group effects, such as response to EW or nest material, were assessed using paired two-tailed Student’s t-tests. Significance threshold was set at P<0.05. The results are shown as mean±SEM. Data processing was performed using TMT Pascal Multi-Target5 software (Framework Computers, Inc., Brighton, MA, USA), and statistically assessed using SPSS V23 (IBM SPSS Statistics, Armonk, NY, USA), or SAS V9.2 (SAS Institute Software Inc., Cary, NC, USA). Figures were prepared using SigmaPlot V12.5 (Systat Software Inc., Chicago, IL, USA), and Adobe Illustrator CC 2015 (Adobe Systems).

## Acknowledgments

A. V. thanks Paul Feinstein and the late Andrea Waltz for advice on gene targeting. We thank Mehdi Tafti for critical insight, Christina Schrick and Yann Emmenegger for superb technical assistance, Arnaud Paradis for help in confocal microscopy, and Andrea Becchetti for comments on the manuscript.

## Author Contributions

A.V. created the *Hcrtr1^flox^* and *Hcrtr1^KO-Gfp^* genomic alleles and mouse strains, designed the project, and wrote the paper; S.L. recorded mice and performed analysis; P.F. elaborated all ECoG analytical methods, performed analysis, and commented on the manuscript.

## Data availability

All materials, data and protocols are available upon request with no restriction.

## References

1 Cirelli, C. The genetic and molecular regulation of sleep: from fruit flies to humans. Nat Rev Neurosci 10, 549–560, doi:10.1038/nrn2683 (2009).

2 Ding, F. et al. Changes in the composition of brain interstitial ions control the sleep-wake cycle. Science 352, 550–555, doi:10.1126/science.aad4821 (2016).

3 Albouy, P., Weiss, A., Baillet, S. & Zatorre, R. J. Selective Entrainment of Theta Oscillations in the Dorsal Stream Causally Enhances Auditory Working Memory Performance. Neuron 94, 193-+, doi:10.1016/j.neuron.2017.03.015 (2017).

4 Schaich Borg, J. et al. Rat intersubjective decisions are encoded by frequency-specific oscillatory contexts. Brain Behav 7, e00710, doi:10.1002/brb3.710 (2017).

5 Trimper, J. B., Galloway, C. R., Jones, A. C., Mandi, K. & Manns, J. R. Gamma Oscillations in Rat Hippocampal Subregions Dentate Gyrus, CA3, CA1, and Subiculum Underlie Associative Memory Encoding. Cell Rep 21, 2419–2432, doi:10.1016/j.celrep.2017.10.123 (2017).

6 Lundqvist, M., Herman, P., Warden, M. R., Brincat, S. L. & Miller, E. K. Gamma and beta bursts during working memory readout suggest roles in its volitional control. Nat Commun 9, 394, doi:10.1038/s41467-017-02791-8 (2018).

7 Blouin, A. M. et al. Human hypocretin and melanin-concentrating hormone levels are linked to emotion and social interaction. Nat Commun 4, 1547, doi:10.1038/ncomms2461 (2013).

8 Giardino, W. J. & de Lecea, L. Hypocretin (orexin) neuromodulation of stress and reward pathways. Curr Opin Neurobiol 29, 103–108, doi:10.1016/j.conb.2014.07.006 (2014).

9 Puskas, N., Papp, R. S., Gallatz, K. & Palkovits, M. Interactions between orexin-immunoreactive fibers and adrenaline or noradrenaline-expressing neurons of the lower brainstem in rats and mice. Peptides 31, 1589–1597, doi:10.1016/j.peptides.2010.04.020 (2010).

10 Schwarz, L. A. & Luo, L. Organization of the locus coeruleus-norepinephrine system. Curr Biol 25, R1051–1056, doi:10.1016/j.cub.2015.09.039 (2015).

11 Berridge, C. W. Noradrenergic modulation of arousal. Brain Research Reviews 58, 1–17, doi:10.1016/j.brainresrev.2007.10.013 (2008).

12 Foote, S. L., Astonjones, G. & Bloom, F. E. Impulse Activity of Locus Coeruleus Neurons in Awake Rats and Monkeys Is a Function of Sensory Stimulation and Arousal. Proceedings of the National Academy of Sciences of the United States of America-Biological Sciences 77, 3033–3037, doi:DOI 10.1073/pnas.77.5.3033 (1980).

13 Florin-Lechner, S. M., Druhan, J. P., Aston-Jones, G. & Valentino, R. J. Enhanced norepinephrine release in prefrontal cortex with burst stimulation of the locus coeruleus. Brain Res 742, 89–97 (1996).

14 Devilbiss, D. M. & Waterhouse, B. D. Phasic and Tonic Patterns of Locus Coeruleus Output Differentially Modulate Sensory Network Function in the Awake Rat. Journal of Neurophysiology 105, 69–87, doi:10.1152/jn.00445.2010 (2011).

15 Peyron, C. et al. Neurons containing hypocretin (orexin) project to multiple neuronal systems. J Neurosci 18, 9996–10015 (1998).

16 Horvath, T. L. et al. Hypocretin (orexin) activation and synaptic innervation of the locus coeruleus noradrenergic system. J Comp Neurol 415, 145–159 (1999).

17 Baldo, B. A., Daniel, R. A., Berridge, C. W. & Kelley, A. E. Overlapping distributions of orexin/hypocretin- and dopamine-beta-hydroxylase immunoreactive fibers in rat brain regions mediating arousal, motivation, and stress. Journal of Comparative Neurology 464, 220–237, doi:10.1002/cne.10783 (2003).

18 Hagan, J. J. et al. Orexin A activates locus coeruleus cell firing and increases arousal in the rat. Proc Natl Acad Sci U S A 96, 10911–10916 (1999).

19 van den Pol, A. N. et al. Hypocretin (orexin) enhances neuron activity and cell synchrony in developing mouse GFP-expressing locus coeruleus. J Physiol 541, 169–185 (2002).

20 Bourgin, P. et al. Hypocretin-1 modulates rapid eye movement sleep through activation of locus coeruleus neurons. Journal of Neuroscience 20, 7760–7765 (2000).

21 Walling, S. G., Nutt, D. J., Lalies, M. D. & Harley, C. W. Orexin-A infusion in the locus ceruleus triggers norepinephrine (NE) release and NE-induced long-term potentiation in the dentate gyrus. Journal of Neuroscience 24, 7421–7426, doi:10.1523/JNEUROSCI.1587-04.2004 (2004).

22 Hirota, K. & Kushikata, T. Central noradrenergic neurones and the mechanism of general anaesthesia. British Journal of Anaesthesia 87, 811–813, doi:Doi 10.1093/Bja/87.6.811 (2001).

23 Tose, R. et al. Interaction between orexinergic neurons and NMDA receptors in the control of locus coeruleus-cerebrocortical noradrenergic activity of the rat. Brain Research 1250, 81–87, doi:10.1016/j.brainres.2008.10.041 (2009).

24 Kukkonen, J. P. & Leonard, C. S. Orexin/hypocretin receptor signalling cascades. Br J Pharmacol 171, 314–331, doi:10.1111/bph.12324 (2014).

25 Trivedi, P., Yu, H., MacNeil, D. J., Van der Ploeg, L. H. T. & Guan, X. M. Distribution of orexin receptor mRNA in the rat brain. Febs Letters 438, 71–75, doi:Doi 10.1016/S0014-5793(98)01266-6 (1998).

26 Marcus, J. N. et al. Differential expression of orexin receptors 1 and 2 in the rat brain. J Comp Neurol 435, 6–25 (2001).

27 Adamantidis, A. R., Zhang, F., Aravanis, A. M., Deisseroth, K. & de Lecea, L. Neural substrates of awakening probed with optogenetic control of hypocretin neurons. Nature 450, 420–424, doi:10.1038/nature06310 (2007).

28 Carter, M. E. et al. Tuning arousal with optogenetic modulation of locus coeruleus neurons. Nat Neurosci 13, 1526–1533, doi:10.1038/nn.2682 (2010).

29 Carter, M. E. et al. Mechanism for Hypocretin-mediated sleep-to-wake transitions. Proc Natl Acad Sci U S A 109, E2635–2644, doi:10.1073/pnas.1202526109 (2012).

30 Chemelli, R. M. et al. Narcolepsy in orexin knockout mice: molecular genetics of sleep regulation. Cell 98, 437–451 (1999).

31 Hara, J. et al. Genetic ablation of orexin neurons in mice results in narcolepsy, hypophagia, and obesity. Neuron 30, 345–354 (2001).

32 Hasegawa, E., Yanagisawa, M., Sakurai, T. & Mieda, M. Orexin neurons suppress narcolepsy via 2 distinct efferent pathways. Journal of Clinical Investigation 124, 604–616, doi:10.1172/JCI71017 (2014).

33 Hunsley, M. S. & Palmiter, R. D. Norepinephrine-deficient mice exhibit normal sleep-wake states but have shorter sleep latency after mild stress and low doses of amphetamine. Sleep 26, 521–526 (2003).

34 Ouyang, M., Hellman, K., Abel, T. & Thomas, S. A. Adrenergic signaling plays a critical role in the maintenance of waking and in the regulation of REM sleep. Journal of Neurophysiology 92, 2071–2082, doi:10.1152/jn.00226.2004 (2004).

35 Singh, C., Oikonomou, G. & Prober, D. A. Norepinephrine is required to promote wakefulness and for hypocretin-induced arousal in zebrafish. Elife 4, doi:10.7554/eLife.07000 (2015).

36 Parlato, R., Otto, C., Begus, Y., Stotz, S. & Schutz, G. Specific ablation of the transcription factor CREB in sympathetic neurons surprisingly protects against developmentally regulated apoptosis. Development 134, 1663–1670, doi:10.1242/dev.02838 (2007).

37 Langer, S. Z. Presynaptic regulation of the release of catecholamines. Pharmacol Rev 32, 337–362 (1980).

38 Xu, Z. Q., Shi, T. J. & Hokfelt, T. Galanin/GMAP- and NPY-like immunoreactivities in locus coeruleus and noradrenergic nerve terminals in the hippocampal formation and cortex with notes on the galanin-R1 and -R2 receptors. J Comp Neurol 392, 227–251 (1998).

39 Schlicker, E. & Kathmann, M. Presynaptic neuropeptide receptors. Handb Exp Pharmacol, 409–434, doi:10.1007/978-3-540-74805-2-13 (2008).

40 Burlet, S., Tyler, C. J. & Leonard, C. S. Direct and indirect excitation of laterodorsal tegmental neurons by Hypocretin/Orexin peptides: implications for wakefulness and narcolepsy. J Neurosci 22, 2862–2872, doi:20026234 (2002).

41 van den Pol, A. N., Gao, X. B., Obrietan, K., Kilduff, T. S. & Belousov, A. B. Presynaptic and postsynaptic actions and modulation of neuroendocrine neurons by a new hypothalamic peptide, Hypocretin/Orexin. Journal of Neuroscience 18, 7962–7971 (1998).

42 Li, Y., Gao, X. B., Sakurai, T. & van den Pol, A. N. Hypocretin/Orexin excites hypocretin neurons via a local glutamate neuron-A potential mechanism for orchestrating the hypothalamic arousal system. Neuron 36, 1169–1181 (2002).

43 Lambe, E. K. & Aghajanian, G. K. Hypocretin (orexin) induces calcium transients in single spines postsynaptic to identified thalamocortical boutons in prefrontal slice. Neuron 40, 139–150 (2003).

44 Belle, M. D. et al. Acute suppressive and long-term phase modulation actions of orexin on the mammalian circadian clock. J Neurosci 34, 3607–3621, doi:10.1523/JNEUROSCI.3388-13.2014 (2014).

45 Aracri, P., Banfi, D., Pasini, M. E., Amadeo, A. & Becchetti, A. Hypocretin (orexin) regulates glutamate input to fast-spiking interneurons in layer V of the Fr2 region of the murine prefrontal cortex. Cereb Cortex 25, 1330–1347, doi:10.1093/cercor/bht326 (2015).

46 Baimel, C. et al. Orexin/hypocretin role in reward: implications for opioid and other addictions. Br J Pharmacol 172, 334–348, doi:10.1111/bph.12639 (2015).

47 Wenger Combremont, A. L., Bayer, L., Dupre, A., Muhlethaler, M. & Serafin, M. Effects of Hypocretin/Orexin and Major Transmitters of Arousal on Fast Spiking Neurons in Mouse Cortical Layer 6B. Cereb Cortex 26, 3553–3562, doi:10.1093/cercor/bhw158 (2016).

48 Li, S., Franken, P. & Vassalli, A. Altered allostatic regulation of wakefulness and slow-wave-sleep spectral quality in mice with hypocretin (orexin) receptor 1 inactivation in noradrenergic cells. bioRxiv (2017).

49 Lakso, M. et al. Efficient in vivo manipulation of mouse genomic sequences at the zygote stage. Proceedings of the National Academy of Sciences of the United States of America 93, 5860–5865, doi:Doi 10.1073/Pnas.93.12.5860 (1996).

50 Vassalli, A., Li, S. & Tafti, M. Did hypocretin receptor 2 autoantibodies cause narcolepsy with hypocretin deficiency in Pandemrix-vaccinated children? Comment on “Antibodies to influenza nucleoprotein cross-react with human hypocretin receptor 2”. Sci Transl Med 7, 314le312, doi:10.1126/scitranslmed.aad2353 (2015).

51 Franken, P., Dijk, D. J., Tobler, I. & Borbely, A. A. Sleep deprivation in rats: effects on EEG power spectra, vigilance states, and cortical temperature. Am J Physiol 261, R198–208 (1991).

52 Cajochen, C., Wyatt, J. K., Czeisler, C. A. & Dijk, D. J. Separation of circadian and wake duration-dependent modulation of EEG activation during wakefulness. Neuroscience 114, 1047–1060, doi:Doi 10.1016/S0306-4522(02)00209-9 (2002).

53 Vassalli, A. & Franken, P. Hypocretin (orexin) is critical in sustaining theta/gamma-rich waking behaviors that drive sleep need. Proc Natl Acad Sci U S A 114, E5464–E5473, doi:10.1073/pnas.1700983114 (2017).

54 Buzsaki, G. Theta oscillations in the hippocampus. Neuron 33, 325–340 (2002).

55 Vyazovskiy, V. V. & Tobler, I. Theta activity in the waking EEG is a marker of sleep propensity in the rat. Brain Res 1050, 64–71, doi:10.1016/j.brainres.2005.05.022 (2005).

56 Vandecasteele, M. et al. Optogenetic activation of septal cholinergic neurons suppresses sharp wave ripples and enhances theta oscillations in the hippocampus. Proc Natl Acad Sci U S A 111, 13535–13540, doi:10.1073/pnas.1411233111 (2014).

57 McFarland, W. L., Teitelbaum, H. & Hedges, E. K. Relationship between hippocampal theta activity and running speed in the rat. J Comp Physiol Psychol 88, 324–328 (1975).

58 Slawinska, U. & Kasicki, S. The frequency of rat’s hippocampal theta rhythm is related to the speed of locomotion. Brain Res 796, 327–331 (1998).

59 Gaskill, B. N., Karas, A. Z., Garner, J. P. & Pritchett-Corning, K. R. Nest Building as an Indicator of Health and Welfare in Laboratory Mice. Jove-Journal of Visualized Experiments, doi:10.3791/51012 (2013).

60 Eban-Rothschild, A., Rothschild, G., Giardino, W. J., Jones, J. R. & de Lecea, L. VTA dopaminergic neurons regulate ethologically relevant sleep-wake behaviors. Nat Neurosci 19, 1356–1366, doi:10.1038/nn.4377 (2016).

61 Crowell, A. L. et al. Oscillations in sensorimotor cortex in movement disorders: an electrocorticography study. Brain 135, 615–630, doi:10.1093/brain/awr332 (2012).

62 Delaville, C., McCoy, A. J., Gerber, C. M., Cruz, A. V. & Walters, J. R. Subthalamic nucleus activity in the awake hemiparkinsonian rat: relationships with motor and cognitive networks. J Neurosci 35, 6918–6930, doi:10.1523/JNEUROSCI.0587-15.2015 (2015).

63 Grey, J. & McNaughton, N. The Neuropshychology of Anxiety: an Enquiry into the Functions of the Septo-Hippocampal System. 2nd Edn edn, (Oxford University Press, 2003).

64 Daan, S., Beersma, D. G. & Borbely, A. A. Timing of human sleep: recovery process gated by a circadian pacemaker. Am J Physiol 246, R161–183 (1984).

65 Franken, P., Chollet, D. & Tafti, M. The homeostatic regulation of sleep need is under genetic control. J Neurosci 21, 2610–2621 (2001).

66 Cirelli, C., Huber, R., Gopalakrishnan, A., Southard, T. L. & Tononi, G. Locus ceruleus control of slow-wave homeostasis. J Neurosci 25, 4503–4511, doi:10.1523/JNEUROSCI.4845-04.2005 (2005).

67 Burgess, N. & O’Keefe, J. Models of place and grid cell firing and theta rhythmicity. Current Opinion in Neurobiology 21, 734–744, doi:10.1016/j.conb.2011.07.002 (2011).

68 Wenger Combremont, A. L., Bayer, L., Dupre, A., Muhlethaler, M. & Serafin, M. Slow Bursting Neurons of Mouse Cortical Layer 6b Are Depolarized by Hypocretin/Orexin and Major Transmitters of Arousal. Front Neurol 7, 88, doi:10.3389/fneur.2016.00088 (2016).

69 Kempadoo, K. A., Mosharov, E. V., Choi, S. J., Sulzer, D. & Kandel, E. R. Dopamine release from the locus coeruleus to the dorsal hippocampus promotes spatial learning and memory. Proceedings of the National Academy of Sciences of the United States of America 113, 14835–14840, doi:10.1073/pnas.1616515114 (2016).

70 Gao, X. B. & Hermes, G. Neural plasticity in hypocretin neurons: the basis of hypocretinergic regulation of physiological and behavioral functions in animals. Front Syst Neurosci 9, 142, doi:10.3389/fnsys.2015.00142 (2015).

71 Valko, P. O. et al. Increase of histaminergic tuberomammillary neurons in narcolepsy. Ann Neurol 74, 794–804, doi:10.1002/ana.24019 (2013).

72 Holloway, B. B. et al. Monosynaptic glutamatergic activation of locus coeruleus and other lower brainstem noradrenergic neurons by the C1 cells in mice. J Neurosci 33, 18792–18805, doi:10.1523/JNEUROSCI.2916-13.2013 (2013).

73 Franken, P., Tobler, I. & Borbely, A. A. Sleep homeostasis in the rat: simulation of the time course of EEG slow-wave activity. Neurosci Lett 130, 141–144 (1991).

74 Dijk, D. J., Duffy, J. F. & Czeisler, C. A. Circadian and Sleep Wake Dependent Aspects of Subjective Alertness and Cognitive Performance. Journal of Sleep Research 1, 112–117 (1992).

75 Cajochen, C., Brunner, D. P., Krauchi, K., Graw, P. & Wirz-Justice, A. Power density in theta/alpha frequencies of the waking EEG progressively increases during sustained wakefulness. Sleep 18, 890–894 (1995).

76 Aeschbach, D. et al. Dynamics of the human EEG during prolonged wakefulness: evidence for frequency-specific circadian and homeostatic influences. Neuroscience Letters 239, 121–124, doi:Doi 10.1016/S0304-3940(97)00904-X (1997).

77 Finelli, L. A., Baumann, H., Borbely, A. A. & Achermann, P. Dual electroencephalogram markers of human sleep homeostasis: correlation between theta activity in waking and slow-wave activity in sleep. Neuroscience 101, 523–529 (2000).

78 Huber, R., Deboer, T. & Tobler, I. Topography of EEG dynamics after sleep deprivation in mice. Journal of Neurophysiology 84, 1888–1893 (2000).

79 Lal, S. K. L. & Craig, A. Driver fatigue: Electroencephalography and psychological assessment. Psychophysiology 39, 313–321, doi:10.1017/S0048577201393095 (2002).

80 Gronli, J., Rempe, M. J., Clegern, W., Schmidt, M. & Wisor, J. P. Beta EEG reflects sensory processing in active wakefulness and homeostatic sleep drive in quiet wakefulness. Journal of Sleep Research 25, 257–268, doi:10.1111/jsr.12380 (2016).

81 Krueger, J. M. et al. Sleep as a fundamental property of neuronal assemblies. Nature Reviews Neuroscience 9, 910–919, doi:10.1038/nrn2521 (2008).

82 Lemaire, N. et al. Effects of dopamine depletion on LFP oscillations in striatum are task- and learning-dependent and selectively reversed by L-DOPA. Proc Natl Acad Sci U S A 109, 18126–18131, doi:10.1073/pnas.1216403109 (2012).

83 Vassalli, A. et al. Electroencephalogram paroxysmal theta characterizes cataplexy in mice and children. Brain 136, 1592–1608, doi:10.1093/brain/awt069 (2013).

84 Brown, R. A., Walling, S. G., Milway, J. S. & Harley, C. W. Locus ceruleus activation suppresses feedforward interneurons and reduces beta-gamma electroencephalogram frequencies while it enhances theta frequencies in rat dentate gyrus. J Neurosci 25, 1985–1991, doi:10.1523/JNEUROSCI.4307-04.2005 (2005).

85 Berridge, C. W. & Foote, S. L. Effects of Locus-Coeruleus Activation on Electroencephalographic Activity in Neocortex and Hippocampus. Journal of Neuroscience 11, 3135–3145 (1991).

86 Walling, S. G., Brown, R. A., Milway, J. S., Earle, A. G. & Harley, C. W. Selective tuning of hippocampal oscillations by phasic locus coeruleus activation in awake male rats. Hippocampus 21, 1250–1262, doi:10.1002/hipo.20816 (2011).

87 Vazey, E. M. & Aston-Jones, G. Designer receptor manipulations reveal a role of the locus coeruleus noradrenergic system in isoflurane general anesthesia. Proc Natl Acad Sci U S A 111, 3859–3864, doi:10.1073/pnas.1310025111 (2014).

88 Sara, S. J. Locus Coeruleus in time with the making of memories. Curr Opin Neurobiol 35, 87–94, doi:10.1016/j.conb.2015.07.004 (2015).

89 Alreja, M. & Liu, W. Noradrenaline induces IPSCs in rat medial septal/diagonal band neurons: involvement of septohippocampal GABAergic neurons. J Physiol 494 (Pt 1), 201–215 (1996).

90 Wu, M. et al. Hypocretin increases impulse flow in the septohippocampal GABAergic pathway: implications for arousal via a mechanism of hippocampal disinhibition. J Neurosci 22, 7754–7765 (2002).

91 Berridge, C. W., Schmeichel, B. E. & Espana, R. A. Noradrenergic modulation of wakefulness/arousal. Sleep Med Rev 16, 187–197, doi:10.1016/j.smrv.2011.12.003 (2012).

92 Gerashchenko, D., Salin-Pascual, R. & Shiromani, P. J. Effects of hypocretin-saporin injections into the medial septum on sleep and hippocampal theta. Brain Res 913, 106–115 (2001).

93 Selbach, O. et al. Orexins/hypocretins cause sharp wave- and theta-related synaptic plasticity in the hippocampus via glutamatergic, gabaergic, noradrenergic, and cholinergic signaling. Neuroscience 127, 519–528, doi:10.1016/j.neuroscience.2004.05.012 (2004).

94 Hangya, B., Borhegyi, Z., Szilagyi, N., Freund, T. F. & Varga, V. GABAergic neurons of the medial septum lead the hippocampal network during theta activity. J Neurosci 29, 8094–8102, doi:10.1523/JNEUROSCI.5665-08.2009 (2009).

95 Amilhon, B. et al. Parvalbumin Interneurons of Hippocampus Tune Population Activity at Theta Frequency. Neuron 86, 1277–1289, doi:10.1016/j.neuron.2015.05.027 (2015).

96 Wulff, P. et al. Hippocampal theta rhythm and its coupling with gamma oscillations require fast inhibition onto parvalbumin-positive interneurons. Proc Natl Acad Sci U S A 106, 3561–3566, doi:10.1073/pnas.0813176106 (2009).

97 Korotkova, T., Fuchs, E. C., Ponomarenko, A., von Engelhardt, J. & Monyer, H. NMDA receptor ablation on parvalbumin-positive interneurons impairs hippocampal synchrony, spatial representations, and working memory. Neuron 68, 557–569, doi:10.1016/j.neuron.2010.09.017 (2010).

98 Varga, C. et al. Functional Fission of Parvalbumin Interneuron Classes During Fast Network Events. Elife 3, doi:10.7554/eLife.04006 (2014).

99 Mather, M., Clewett, D., Sakaki, M. & Harley, C. W. Norepinephrine ignites local hot spots of neuronal excitation: How arousal amplifies selectivity in perception and memory. Behav Brain Sci, 1–100, doi:10.1017/S0140525X15000667 (2015).

100 Cape, E. G. & Jones, B. E. Differential modulation of high-frequency gamma-electroencephalogram activity and sleep-wake state by noradrenaline and serotonin microinjections into the region of cholinergic basalis neurons. J Neurosci 18, 2653–2666 (1998).

101 Bisetti, A. et al. Excitatory action of hypocretin/orexin on neurons of the central medial amygdala. Neuroscience 142, 999–1004, doi:10.1016/j.neuroscience.2006.07.018 (2006).

102 McCall, J. G. et al. Locus coeruleus to basolateral amygdala noradrenergic projections promote anxiety-like behavior. Elife 6, doi:10.7554/eLife.2017.e18247 (2017).

103 Mejias-Aponte, C. A., Drouin, C. & Aston-Jones, G. Adrenergic and noradrenergic innervation of the midbrain ventral tegmental area and retrorubral field: prominent inputs from medullary homeostatic centers. J Neurosci 29, 3613–3626, doi:10.1523/JNEUROSCI.4632-08.2009 (2009).

104 Sears, R. M. et al. Orexin/hypocretin system modulates amygdala-dependent threat learning through the locus coeruleus. Proc Natl Acad Sci U S A 110, 20260–20265, doi:10.1073/pnas.1320325110 (2013).

105 Flores, A., Herry, C., Maldonado, R. & Berrendero, F. Facilitation of Contextual Fear Extinction by Orexin-1 Receptor Antagonism Is Associated with the Activation of Specific Amygdala Cell Subpopulations. International Journal of Neuropsychopharmacology 20, 654–659, doi:10.1093/ijnp/pyx029 (2017).

106 Soya, S. et al. Orexin modulates behavioral fear expression through the locus coeruleus. Nature Communications 8, doi:10.1038/S41467-017-01782-Z (2017).

107 Flores, A. et al. The Hypocretin/Orexin System Mediates the Extinction of Fear Memories. Neuropsychopharmacology 39, 2732–2741, doi:10.1038/npp.2014.146 (2014).

108 Ponz, A. et al. Abnormal activity in reward brain circuits in human narcolepsy with cataplexy. Ann Neurol 67, 190–200, doi:10.1002/ana.21825 (2010).

109 Bayard, S. et al. Decision making in narcolepsy with cataplexy. Sleep 34, 99–104 (2011).

110 Dimitrova, A. et al. Reward-seeking behavior in human narcolepsy. J Clin Sleep Med 7, 293–300, doi:10.5664/JCSM.1076 (2011).

111 Merlotti, E. et al. Impulsiveness in patients with bulimia nervosa: electrophysiological evidence of reduced inhibitory control. Neuropsychobiology 68, 116–123, doi:10.1159/000352016 (2013).

112 Chen, X. W. et al. Hypocretin-1 potentiates NMDA receptor-mediated somatodendritic secretion from locus ceruleus neurons. J Neurosci 28, 3202–3208, doi:10.1523/JNEUROSCI.4426-07.2008 (2008).

113 Agnati, L. F., Guidolin, D., Guescini, M., Genedani, S. & Fuxe, K. Understanding wiring and volume transmission. Brain Res Rev 64, 137–159, doi:10.1016/j.brainresrev.2010.03.003 (2010).

114 Arima, J., Kubo, C., Ishibashi, H. & Akaike, N. alpha2-Adrenoceptor-mediated potassium currents in acutely dissociated rat locus coeruleus neurones. J Physiol 508 (Pt 1), 57–66 (1998).

115 Rasmussen, K. The role of the locus coeruleus and N-methylD-aspartic acid (NMDA) and AMPA receptors in opiate withdrawal. Neuropsychopharmacology 13, 295–300, doi:10.1016/0893-133X(95)00082-O (1995).

116 Huang, H. P. et al. Long latency of evoked quantal transmitter release from somata of locus coeruleus neurons in rat pontine slices. Proceedings of the National Academy of Sciences of the United States of America 104, 1401–1406, doi:10.1073/pnas.0608897104 (2007).

117 Nishino, S. & Mignot, E. Pharmacological aspects of human and canine narcolepsy. Progress in Neurobiology 52, 27–78, doi:Doi 10.1016/S0301-0082(96)00070-6 (1997).

118 Wu, M. F. et al. Locus coeruleus neurons: Cessation of activity during cataplexy. Neuroscience 91, 1389–1399, doi:Doi 10.1016/S0306-4522(98)00600-9 (1999).

119 Burgess, C. R. & Peever, J. H. A noradrenergic mechanism functions to couple motor behavior with arousal state. Curr Biol 23, 1719–1725, doi:10.1016/j.cub.2013.07.014 (2013).

120 Hasegawa, E. et al. Serotonin neurons in the dorsal raphe mediate the anticataplectic action of orexin neurons by reducing amygdala activity. Proc Natl Acad Sci U S A 114, E3526–E3535, doi:10.1073/pnas.1614552114 (2017).

121 Huber, R., Ghilardi, M. F., Massimini, M. & Tononi, G. Local sleep and learning. Nature 430, 78–81, doi:10.1038/nature02663 (2004).

122 Steriade, M., Contreras, D., Curro Dossi, R. & Nunez, A. The slow (< 1 hz) oscillation in reticular thalamic and thalamocortical neurons: scenario of sleep rhythm generation in interacting thalamic and neocortical networks. J Neurosci 13, 3284–3299 (1993).

123 Achermann, P. & Borbely, A. A. Low-frequency (< 1 hz) oscillations in the human sleep electroencephalogram. Neuroscience 81, 213–222 (1997).

124 Bersagliere, A., Pascual-Marqui, R. D., Tarokh, L. & Achermann, P. Mapping Slow Waves by EEG Topography and Source Localization: Effects of Sleep Deprivation. Brain Topogr, doi:10.1007/s10548-017-0595-6 (2017).

125 Mallick, B. N. et al. Role of norepinephrine in the regulation of rapid eye movement sleep. J Biosci 27, 539–551 (2002).

126 Dauvilliers, Y., Arnulf, I. & Mignot, E. Narcolepsy with cataplexy. Lancet 369, 499–511, doi:10.1016/S0140-6736(07)60237-2 (2007).

127 Roman, A., Meftah, S., Arthaud, S., Luppi, P. H. & Peyron, C. The Inappropriate Occurrence of REM Sleep in Narcolepsy is not due to a Defect in Homeostatic Regulation of REM Sleep. Sleep, doi:10.1093/sleep/zsy046 (2018).

128 Mochizuki, T. et al. Behavioral state instability in orexin knock-out mice. J Neurosci 24, 6291–6300, doi:10.1523/JNEUROSCI.0586-04.2004 (2004).

129 Hoyer, D. & Jacobson, L. H. Orexin in sleep, addiction and more: Is the perfect insomnia drug at hand? Neuropeptides 47, 477–488, doi:10.1016/j.npep.2013.10.009 (2013).

130 Mather, M. & Harley, C. W. The Locus Coeruleus: Essential for Maintaining Cognitive Function and the Aging Brain. Trends Cogn Sci 20, 214–226, doi:10.1016/j.tics.2016.01.001 (2016).

131 Chen, J. & Randeva, H. S. Genomic organization of mouse orexin receptors: characterization of two novel tissue-specific splice variants. Mol Endocrinol 18, 2790–2804, doi:10.1210/me.2004-0167 (2004).

132 Rodriguez, C. I. et al. High-efficiency deleter mice show that FLPe is an alternative to Cre-loxP. Nature Genetics 25, 139–140 (2000).

133 Mang, G. M. & Franken, P. Sleep and EEG Phenotyping in Mice. Curr Protoc Mouse Biol 2, 55–74, doi:10.1002/9780470942390.mo110126 (2012).

134 Franken, P., Malafosse, A. & Tafti, M. Genetic variation in EEG activity during sleep in inbred mice. Am J Physiol 275, R1127–1137 (1998).

135 Hasan, S. et al. How to keep the brain awake? The complex molecular pharmacogenetics of wake promotion. Neuropsychopharmacology 34, 1625–1640, doi:10.1038/npp.2009.3 (2009).

136 Franklin, K. B. J. & Paxinos, G. Paxinos and Franklin’s The mouse brain in stereotaxic coordinates. Fourth edition. edn.

